# Notch signalling plays a role in patterning the ventral mesoderm during early embryogenesis in *Drosophila melanogaster*

**DOI:** 10.1101/2023.09.27.558900

**Authors:** Marvel Megaly, Gregory Foran, Arsala Ali, Anel Turgambayeva, Ryan D. Hallam, Ping Liang, Aleksandar Necakov

**Affiliations:** Biological Sciences Department, Brock University, St. Catharines, Ontario, Canada; Neuroscience Department, Baylor College of Medicine, Houston, Texas, United States

**Keywords:** keyword 1, keyword 2, keyword 3 (List three to ten pertinent keywords specific to the article yet reasonably common within the subject discipline.)

## Abstract

Notch signalling is a critical regulator of multiple developmental processes through its ability to control gene expression and thereby influence cell fate specification and cell proliferation through direct cell-cell communication. Although Notch signalling has been implicated in myogenesis during late embryogenesis, its role in early mesoderm development has been largely unexplored. Endocytosis of the Notch ligand Delta and the Notch receptor extracellular domain, a critical step in Notch pathway activation, has been extensively observed in the ventral mesoderm of the early *Drosophila* embryo, indicating a potential for Notch signalling activity in this early germ layer. Here we present evidence that genes critical to mesoderm development require and are responsive to Notch signalling activity. Using a novel light-inducible Optogenetic variant of the Notch intracellular domain (OptoNotch), which affords precise spatial and temporal control over ectopic activation of Notch signalling, in combination with high-resolution fluorescent RNA *in situ* hybridization and qPCR, we identified a set of mesodermal genes whose expression is directly regulated by Notch signalling. We also provide evidence that Notch signalling indirectly regulates the dorsal-ventral patterning program mediated by the Toll signalling pathway through the Dorsal/ Twist/ Snail gene network. Our findings demonstrate that Notch signalling regulates ventral mesoderm patterning and is critical for establishing the mesoderm-mesectoderm-ectoderm boundary by regulating gene expression patterns and providing negative feedback on the upstream patterning network.

## 1. Introduction

Embryogenesis is a critical developmental process that is driven by patterns of gene expression, which collectively organize the coordinated cell proliferation and differentiation required for proper organismal development ^[1]^. *Drosophila melanogaster* has been used as a model organism to study the critical mechanisms that direct embryogenesis^[1]^ and has also served as a useful model system to study signalling pathways that coordinate body patterning and morphogenetic movements in the formation of tissues and organs, which collectively give rise to entire organisms ^[1,2]^.

Embryonic development can be divided into four main stages; cleavage, blastulation, gastrulation, and neurulation ^[3]^. The first stage, cleavage, involves multiple rounds of cell division that drive the formation of the blastula from a single-cell zygote ^[3]^. Blastulation subsequently results in the formation of the blastula through subsequent rounds of cell division, with some accompanying differentiation occurring at the end of blastulation^[3]^. Gastrulation involves the development of the three germ layers; the ectoderm, the mesoderm, and the endoderm^2^. The ectoderm forms the outermost layers of the organism; the surface ectoderm, neural crest, and the neural tube^[4]^.The mesoderm forms the mid-layer of the organism, which later develops into muscle, bone, cartilage, the circulatory and lymphatic systems, connective and adipose tissue, the notochord, dermis, serous membranes, and the genitourinary system^[4]^. Finally, the last stage of embryogenesis, neurulation, involves the development of the nervous system^[4]^. During early embryogenesis in *Drosophila melanogaster*, 13 nuclear division cycles occur within a shared syncytial cytoplasm^[3]^. This early developmental stage is instructed by maternally deposited mRNAs and proteins that organize the anterior-posterior and dorsal-ventral body axes, and define the segmental pattern of the embryo that later gives rise to different body segments of the larva and adult^[5]^. This early body plan is subsequently refined by functional products from the zygotic genome^[5]^. During the 14^th^ nuclear cycle, individual cell membranes form to separate, encapsulate, and thus individuate nuclei during blastoderm cellularization^[3]^. Many maternally deposited materials are degraded during this stage, and the control of development becomes increasingly dependent upon zygotic gene expression^[6,7]^. One of the critical factors for establishing the dorsal-ventral axis is a maternally deposited transcription factor, Dorsal, which functions downstream of the Toll signalling pathway and forms a morphogen gradient across the Dorsal-Ventral axis^[8]^. Once Toll signalling is activated, Dorsal translocates to the nucleus, where it activates the expression of master regulators of gastrulation; Twist and Snail, which are also transcription factors that activate genes necessary for mesoderm specification^[9,10]^. At the ventral side of the blastoderm, a 21-cell wide anterior-posterior strip of cells in the ventral-most region of the cellularized blastoderm make up the presumptive ventral mesoderm, which undergo a series of cell shape changes; apical flattening, apical constriction, shortening along the apicobasal axis, followed by cell depolarization, dispersion and migration along the ectoderm, to which they attach^[2,4]^. Cell shortening along the apicobasal axis contributes to ventral furrow invagination and mesoderm ingression^[2]^. Upon completion of mesoderm ingression, FGF signalling between mesodermal cells and the underlying ectoderm facilitates mesoderm collapse against the ectodermal cell layer^[11]^. These mesodermal cells subsequently lose polarity, transition to a mesenchymal cell fate, and undergo two rounds of cell division^[11]^. Following this, FGF signalling facilitates cell migration and spreading along the ectoderm to form a cellular monolayer^[11]^.

Notch signalling is critical to cell proliferation, differentiation, and the development of virtually all tissues that give rise to the adult organism^[1]^. It is also essential to the development of both vertebrates and invertebrates^[12]^. For example, it has been previously shown that *Drosophila melanogaster* embryos homozygous for loss-of-function Notch alleles do not reach adulthood^[13]^ as they exhibit neuralized defects and die during embryogenesis. The Notch receptor, a transmembrane protein, is activated by the binding of its ligand, Delta (Dl), on neighbouring signal-sending cells, followed by ligand internalization and Notch receptor activation through transcytosis of the labile Notch extracellular domain, and subsequent intramembrane cleavage by the gamma secretase complex^[1]^. Notch cleavage triggers the release of the Notch intracellular domain (NICD) from the plasma membrane and its concomitant transport to the nucleus where it regulates gene expression through removal of transcriptional co-repressors bound to suppressor of hairless (Su(H)) and the subsequent recruitment of transcriptional co-activators that cooperatively activate transcription of Notch target genes^[1]^. Ligand internalization through endocytosis has been previously shown to be necessary for Notch activation^[1]^. Intriguingly, although no role for Notch signalling has been established in ventral mesoderm patterning to date, endocytosis of Delta and the extracellular domain of the Notch receptor is observed in all mesodermal cells (Figure 1), indicating that Notch signalling may be operative in ventral mesodermal cells^[14]^. Although the role of Notch signalling during embryogenesis has been extensively studied, little is known about its role in the development of the ventral mesoderm^[1]^. Likewise, the identity of potential Notch target genes in the ventral mesoderm is not yet known.

**Figure 1.**
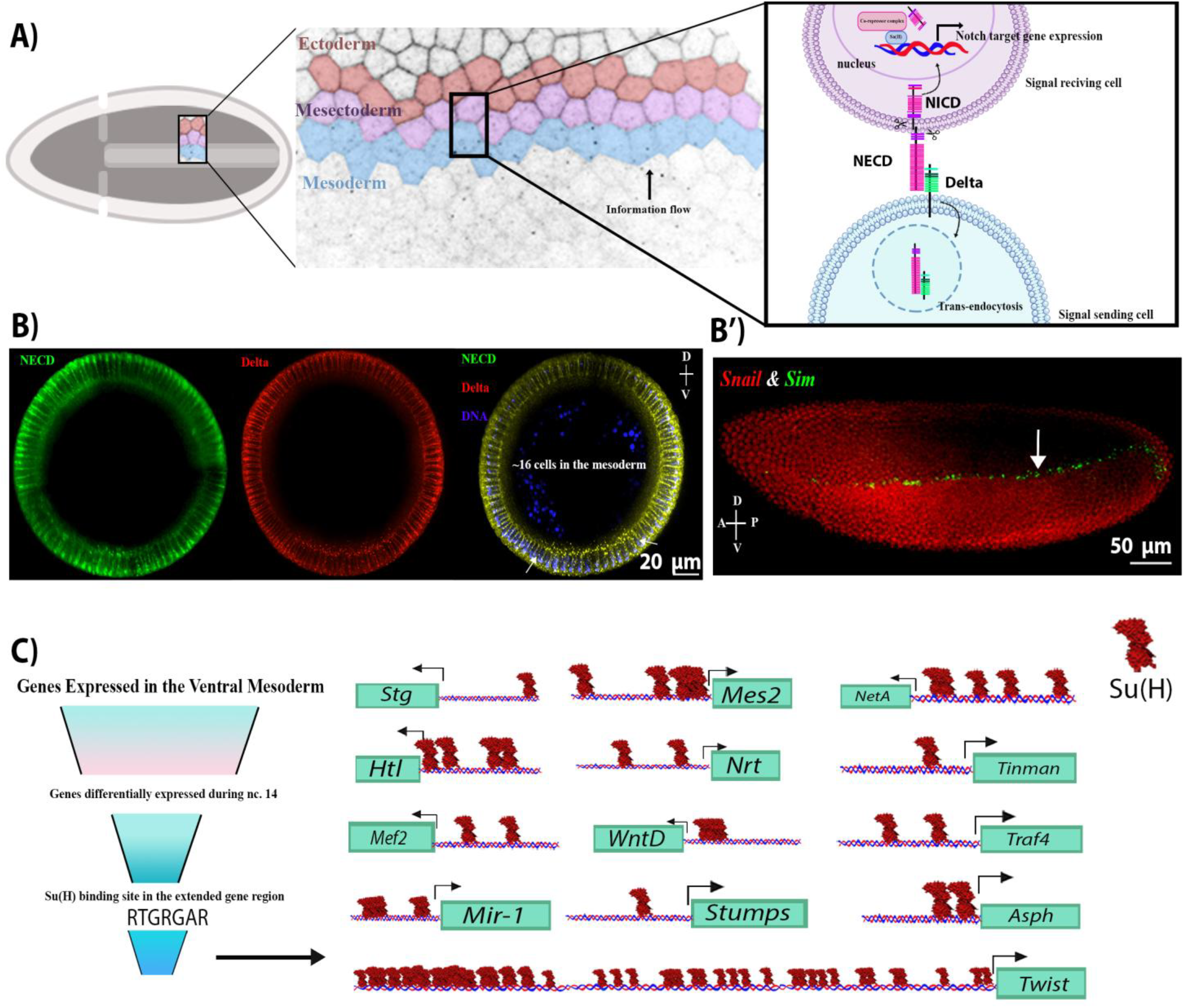
NECD and Delta endocytosis in the mesoderm indicate that Notch signalling is active in the mesoderm. A) a schematic of Notch signalling activation during early embryogenesis in the presumptive ventral mesoderm, neighbouring mesectoderm and ectoderm cells in cellularized embryos. Notch signalling is activated via the binding of the ligand (located on the outermost mesoderm cell), Delta, to the extracellular domain of the Notch receptor (NECD) on a neighbouring mesectoderm cell (signal receiving cell). This interaction leads to a series of cleavage events of the Notch receptor, separating the extracellular and intracellular domains of Notch. NECD and Delta are endocytosed into the signal-sending mesoderm cell, while the intracellular domain of Notch (NICD) is liberated from the plasma membrane and translocates to the nucleus, where it regulates gene expression. B) Cross sections of different wild-type embryos at stage 4 (cellularized blastoderm) showing the internalized Delta and NECD in mesodermal and mesoectodermal (indicated by arrows) cells, indicating endocytosis and therefore Notch signalling activation. B’) A lateral view of stage 4 wild-type embryo showing the RNA expression of a Notch target gene, *Sim* (green), and its repressor, *Snail* (red) in the mesectoderm and mesoderm, respectively. C) Summary of *in silico* analysis showing candidate Notch target genes that were differentially expressed in the mesoderm at nuclear cycle 14 and contained a Su(H) binding site in their proximal promoter. These include; *Asph, Tinman, Traf4, Twist, WntD, Stumps, String Heartless, NetrinA, Neurotactin, mir-1, Mes2*, and *Mef2*. Only the binding sites preceding the 5’UTR are shown here. Su(H) sites are found upstream of the 5’UTR of all transcripts of the genes above except for *Asph*, and *Twist*.

Here we show that Notch signalling is active in the presumptive mesoderm and is required for proper progression through gastrulation. We identified a set of Notch target genes that are expressed in the ventral mesoderm and characterized their expression in embryos with either Notch pathway loss-of-function or gain-of-function mutations. In addition, we developed and characterized a novel tool, OptoNotch, that provides spatially and temporally precise light-gated activation of Notch signalling. Our findings support a mechanism whereby Notch signalling provides negative feedback control over the early patterning network established by the Toll pathway and thereby affects the Dorsal/ Twist/ Snail signalling network. Taken together, our results provide evidence that Notch signalling plays a critical role in the formation of the ventral mesoderm and in defining and positioning the boundary separating it from the ectoderm during early embryogenesis.

## 2. Results

### 2.1. Delta endocytosis and NECD trans-endocytosis and identification of candidate Notch target genes in the mesoderm

The role of Notch signalling during embryogenesis has been well characterized in the lateral ectoderm and the mesectoderm^[15]^. In the lateral ectoderm, Notch signalling works in concert with EGF and Toll signalling to coordinate neurogenesis^[13,16]^. In the mesectoderm, Notch signalling is required to form the boundary between the ectoderm and mesoderm through transcriptional activation of target genes including *single minded* (*sim*)^[17,18]^. Several previous studies have suggested that Notch signalling is active in the ventral mesoderm, a hypothesis that is supported by the observation that the *Drosophila* Notch extracellular domain (NECD) is endocytosed in a Delta-dependent manner in the ventral mesoderm (Figure 1B)^[14,19,20]^. Specifically, internalization of Delta and concomitant endocytosis of the NECD observed in ventral mesodermal cells (Figure 1B) suggests that Notch signalling may be operant in this tissue^[14]^. Traditionally, mesectodermal expression of *sim, m8, bobble, bearded*, and enhancer of split-related transgenes have been used as markers of Notch signalling in the early embryo^[18,21,22]^, where Delta endocytosis in the ventral mesoderm drives Notch activity in the mesectoderm, representing an intercellular and inter-cell-type flow of information (Figure 1A). However, since both *sim* and *m8* are actively repressed in the ventral mesoderm by Snail; a transcriptional repressor that coordinates mesoderm formation, the activity of Notch signalling in the ventral mesoderm may have been overlooked^[18]^. When Snail repression is removed, as is the case in *snail*^*-*^ mutants, *sim* RNA is detected in the mesoderm^[16]^. Thus, this study aimed to investigate the role of Notch signalling in the ventral mesoderm.

To address this question, we first generated a list of candidate transcriptional targets of Notch signalling using bioinformatic analysis of previously published suppressor of hairless CHIP-seq data to identify genes that contain Su(H) binding sites proximal to their promoter and that exhibit differential expression during nuclear cycle 14 (nc 14), which coincides with the increase of Notch signalling activity. This was performed in conjunction with available RNA *in situ* expression data from the Berkeley *Drosophila* genome project, modENCODE RNA-seq expression data^[17,23,24]^. Using this approach, we identified *Asph, Twist, Tinman, Traf4, WntD, Stumps, String, Heartless, NetrinA, Neurotactin, mir-1, Mes2*, and *Mef2* as candidate transcriptional targets of the Notch signalling pathway; (Figure 1C).

### 2.2. Notch signalling is required for the expression of critical mesodermal genes

To test the necessity of Notch signalling in ventral mesodermal development, we employed temperature-sensitive loss-of-function Delta mutant alleles to analyze the expression of the twelve candidate mesodermal Notch target genes we identified through *in silico* analysis. To accomplish this, we precisely quantified changes in transcript abundance for each of the genes by using a combination of two orthogonal methods, fluorescent RNA *in situ* hybridization (FISH) and quantitative real-time PCR (qRT-PCR).

Through this approach, we identified three classes of genes within this set of candidates based on changes in their expression level in response to the loss of Notch signalling (Figure 2):

Class 1 - Genes whose expression requires Notch signalling. These include: *Asph, Mef2, Mes2, Neurotactin, String, Stumps, Tinman, Traf4, Twist* (Figure 2A, 2D),

Class 2 - Genes whose expression is repressed by Notch signalling; *Heartless* and *WntD* (Figure 2B, 2E)

Class 3 - Genes whose expression is independent of Notch signalling, *NetrinA* (Figure 2C, 2F)

**Figure 2.**
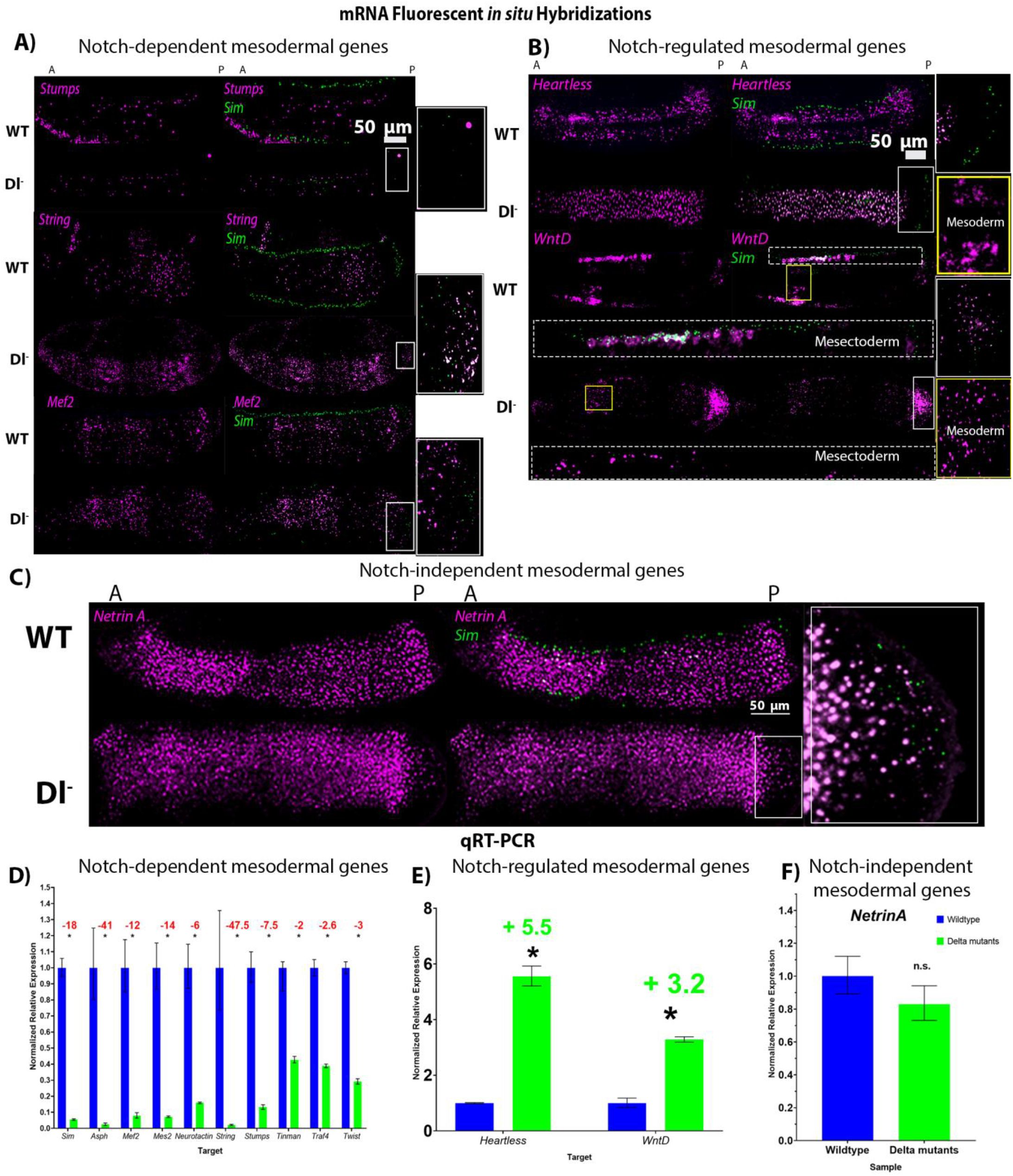
Expression of mesodermal genes in wild-type and loss-of-function Delta mutant embryos show that Notch signalling is necessary for the expression of many mesodermal genes. A-C) fluorescent mRNA *in situ* hybridizations against *sim* (green)and mesodermal genes (purple) in wild-type and Delta mutant embryos at cephalic furrow (stage 5-6). A) shows examples of genes whose expression was significantly reduced in Delta mutants (Dl^-^) compared to their wild-type (WT) counterparts (*Stumps, String*, and *Mef2*). B) shows expression of the two genes whose expression was upregulated in Dl^-^; *Heartless* and *WntD*. C) Expression of *NetrinA* was unchanged in Dl^-^. N= 5 for each genotype per gene. D) qRT-PCR generated normalized relative expression of mesodermal genes and *sim* in wild-type and Delta mutant embryos at cephalic furrow (stage 5-6). Expression of *Asph* (p=0.0007), *Mef2* (p= 0.0007), *Mes2* (p= 7.77419×10^-05^), *Neurotactin* (p=0.0002), *String* (p=0.0004), *Stumps* (p=0.000149162), *Tinman* (p=0.0067), *Traf4* (p=8.19966×10^-05^), and *Twist* (p=5.80546×10^-05^) is significantly reduced in Delta mutants compared to stage-matched wild-type embryos. E) Expression of *Heartless* (p= 1.39725×10^-05^) and *WntD* (p=0.0022) is significantly increased in Delta mutants, while F) the expression of *NetrinA* (p= 0.3359) was not significantly changed. Independent two-tailed T-tests were performed for each gene to compare expression of each gene in Dl^-^ to that in WT embryos. ‘*’ indicates statistical significance determined by a p< 0.05. Error bars represent the standard error of the mean. For each genotype, 30 embryos were used (N=30).

All but one gene (*NetrinA*) showed significant changes in expression pattern between wild-type and Delta mutant embryos. However, *WntD* was expressed at higher levels in the mesoderm of Delta mutants, while significantly reduced in mesectodermal cells, both compared to wild-type embryos (Figure 2B).

### 2.3. Development of OptoNotch allows titration of ectopic Notch signalling activity and incremental expansion of *Sim* expression

The above findings motivated us to test the sufficiency of Notch signalling to drive the expression of the ventral mesoderm target gene candidates and thus we developed a tool that allows us to ectopically activate Notch signalling with precise temporal control. We engineered a light-inducible variant of the NICD (OptoNotch) and expressed it in the early embryo. In this system, prior to photoactivation with blue light (488 nm), OptoNotch is tethered to the membrane. Upon photoactivation, NICD is cleaved through light-inducible reconstitution of TEV protease activity, promoting its translocation to the nucleus where it can activate target gene expression (Figure 3A). We first tested OptoNotch in S2 cells. Prior to photo activation, OptoNotch (tagged with GFP) was localized to the plasma membrane and internal membranous structures but was excluded from the nucleus. As expected, light exposure promotes nuclear translocation of the NotchICD as following one hour of continuous photoactivation, OptoNotch::GFP was detected in the nucleus (Figure 3B).

**Figure 3.**
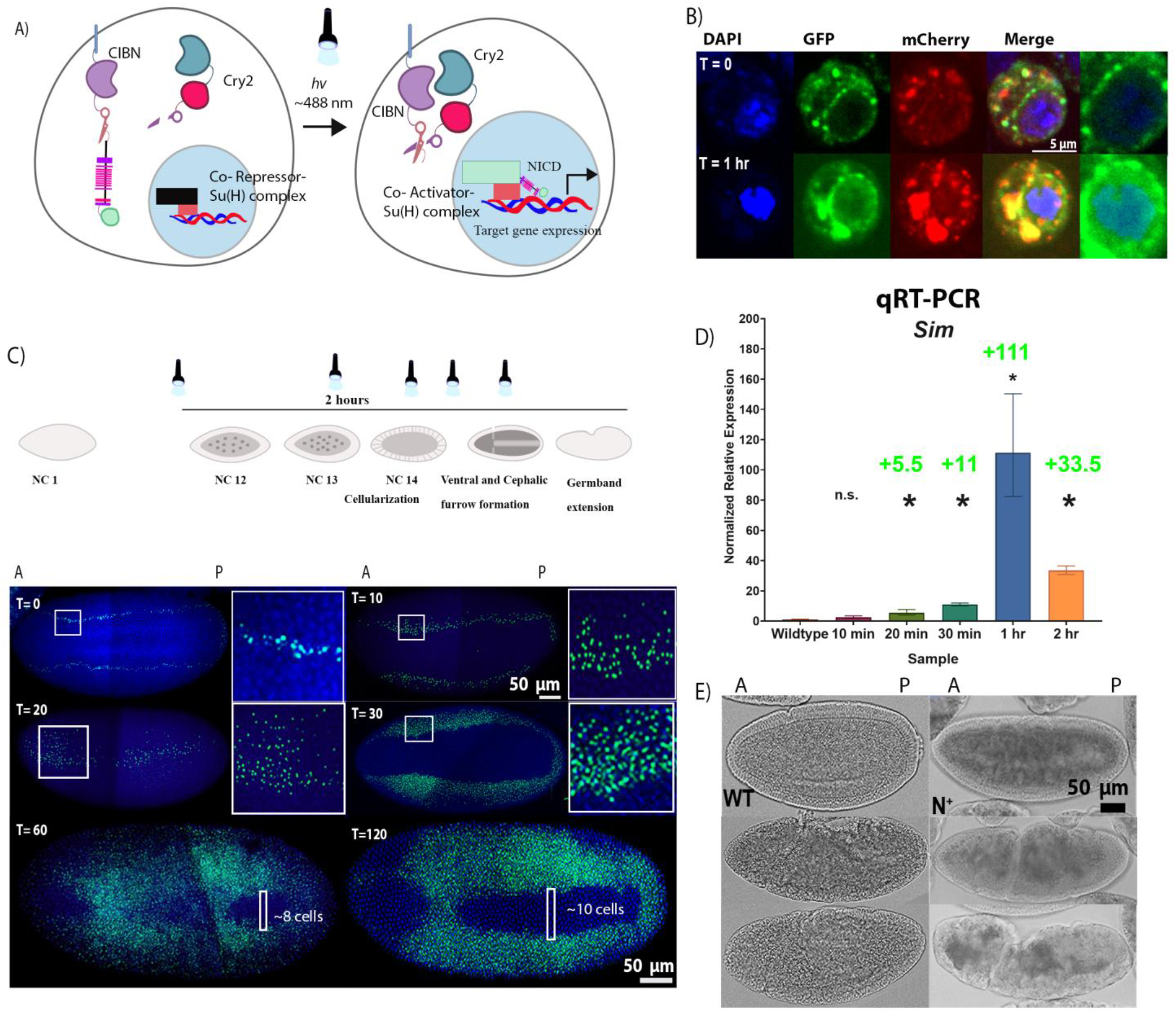
Development of Optogenetic NICD (OptoNotch) revealed competency of mesodermal cells to express a Notch target gene, *sim*. A) Schematic of the mechanism of the Optogenetic variant of NICD (OptoNotch) that we developed which is described in detail in the text. B) S2 cells were transiently transfected with OptoNotch and imaged prior to photoactivation and after continuous photoactivation for 1 hour. Prior to photoactivation (T=0), NICD-GFP is excluded from the nucleus, while after photoactivation. NICD-GFP is observed in the nucleus (blue) after 1 hour (T=1 hr) of continuous photoactivation. The two right-most panels show nuclear GFP at T=0 and T=1 hr. C) Embryos expressing OptoNotch were either not photoactivated (T=0), or continuously photoactivated for 10 (n=3), 20 (n=3), 30 (n=10), 60 (n=10), or 120 (n=30) minutes before fixation. Embryos photoactivated for longer durations were photoactivated starting at an earlier developmental stage (top). Bottom: the corresponding fluorescent *in situ* hybridizations against *sim* (green) in embryos expressing OptoNotch photoactivated for different durations. Nuclei are shown by DAPI nuclear counterstain (blue). Prior to photoactivation (T=0), embryos *sim* is expressed as it is in wild-type embryos (n=10). *Sim* expression is incrementally expanded into lateral regions of OptoNotch embryos photoactivated for 10, 20 and 30 minutes (stage 4 embryos shown, prior to cephalic furrow formation). The regions of *sim* expression expansion are highlighted with white boxes T10 and T20. The adjacent insets were taken from the regions in the white boxes. *Sim* expression is observed in the regions of mesoderm of OptoNotch embryos photoactivated for 60 and 120 minutes (stage 6 embryos shown). In embryos photoactivated for 60 minutes, the maximum number of cells (across the dorsoventral axis) lacking *sim* expression is ∼8 cells, and in those photoactivated for 2 hours, ∼10 cells across the dorsal-ventral axis lack *sim* expression. These regions are highlighted with white boxes in the bottom two images. D) qRT-PCR generated normalized relative expression of *sim* of stage 6 wild-type and OptoNotch embryos photoactivated for different durations, as indicated on the x-axis. N=30 embryos for each group. A one-way ANOVA was performed, p=0.0001, F= 47.68. A post-hoc Tukey HSD test was used to perform pairwise comparisons. For simplicity, the results of comparisons of each group to wild-type embryos are shown, where ‘*’ indicates statistical significance determined by a Tukey’s HSD p<0.05 for that group and WT. “n.s” denotes a p>0.05 between that group and WT. Error bars represent the standard error of the mean. E) Brightfield images of wild-type (WT) and OptoNotch (N+) embryos (photoactivated for 2 hours) at different stages of gastrulation. Morphological defects along the anterior-posterior axis can be observed in N^+^ embryos during early gastrulation and complete breakdown is observed during later stages, N=30 for each genotype.

Next, we expressed our construct in the early embryo. We used *sim* expression as a readout for Notch signalling levels because it is a direct target of Notch signalling. Its expression domain has previously been shown to expand beyond the mesectoderm as a result of overexpression of the NICD^[18]^. As predicted, prior to photoactivation, Notch signalling activity is at similar levels to wild-type embryos, as only endogenous Notch signalling is active (Figure 3C). We also show that we can control the extent of Notch activation by varying the duration of photoactivation (Figure 3C, 3D). Embryos exposed to 10 minutes of light exhibited the smallest expansion in *sim* expression (only seen in mid nc 14 embryos *in situ* but not in older embryos with a cephalic furrow-represented in the qRT-PCR data) (Figure 3C, 3D). Incremental expansions were observed in *sim* expression that correlated with the duration of photoactivation, except that *sim* expression was higher in embryos photoactivated for 1 hour than those photoactivated for 2 hours (Figure 3C, 3D). This could be due to negative feedback mechanisms, which downregulate *sim* expression. After 1 hour of light exposure, the embryos expressed *sim* in the presumptive mesoderm and the number of cells that did not contain *sim* mRNA was decreased significantly compared to wild-type embryos (∼ 8 cells at the most, 0 in some areas along the AP axis). This was also seen in embryos exposed to light for 2 hrs but to a lesser extent (∼ 10 cells at the most, 0 around the cephalic furrow) (Figure 3C, 3D). These results demonstrate the successful developoment and application of a light-inducible NICD variant that provides titratable control over Notch signalling activity, which is sufficient to drive expression of the Notch target gene *sim* in the mesoderm without the removal of the repressor, Snail.

We also observed morphological defects in embryos exposed to light for 1 (not shown) and 2 hours (Figure 3E). These embryos (n=30) exhibited ectopic invaginations along the AP axis during the early stages of gastrulation, where the ventral furrow was forming but the mesoderm was not fully internalized. Following internalization of the mesoderm, the cephalic furrow appears closer to the middle of the mutant embryo along the AP axis compared to wild-type counterparts. There were also additional invaginations along the AP axis on the ventral side. At older stages that roughly correspond to germband retraction, the mutant embryo was twisted, which was reminiscent of the phenotype of loss-of-function Twist mutant embryos (Figure S1).

### 2.4. Ectopic Notch signalling interferes with mesoderm formation and decreases mesodermal target gene expression

To test whether Notch signalling activation is sufficient to drive the expression of the mesodermal genes, we activated Notch signalling using OptoNotch in the early embryo and examined mesodermal gene expression using FISH and qRT-PCR. We photoactivated nc 9-10 embryos expressing both components of the system by exposing them to light, using an LED light box, for 2 hours continuously.

We found that there are two classes of genes within this set of candidates based on their expression level when Notch signalling is overactivated (Figure 4):

**Figure 4.**
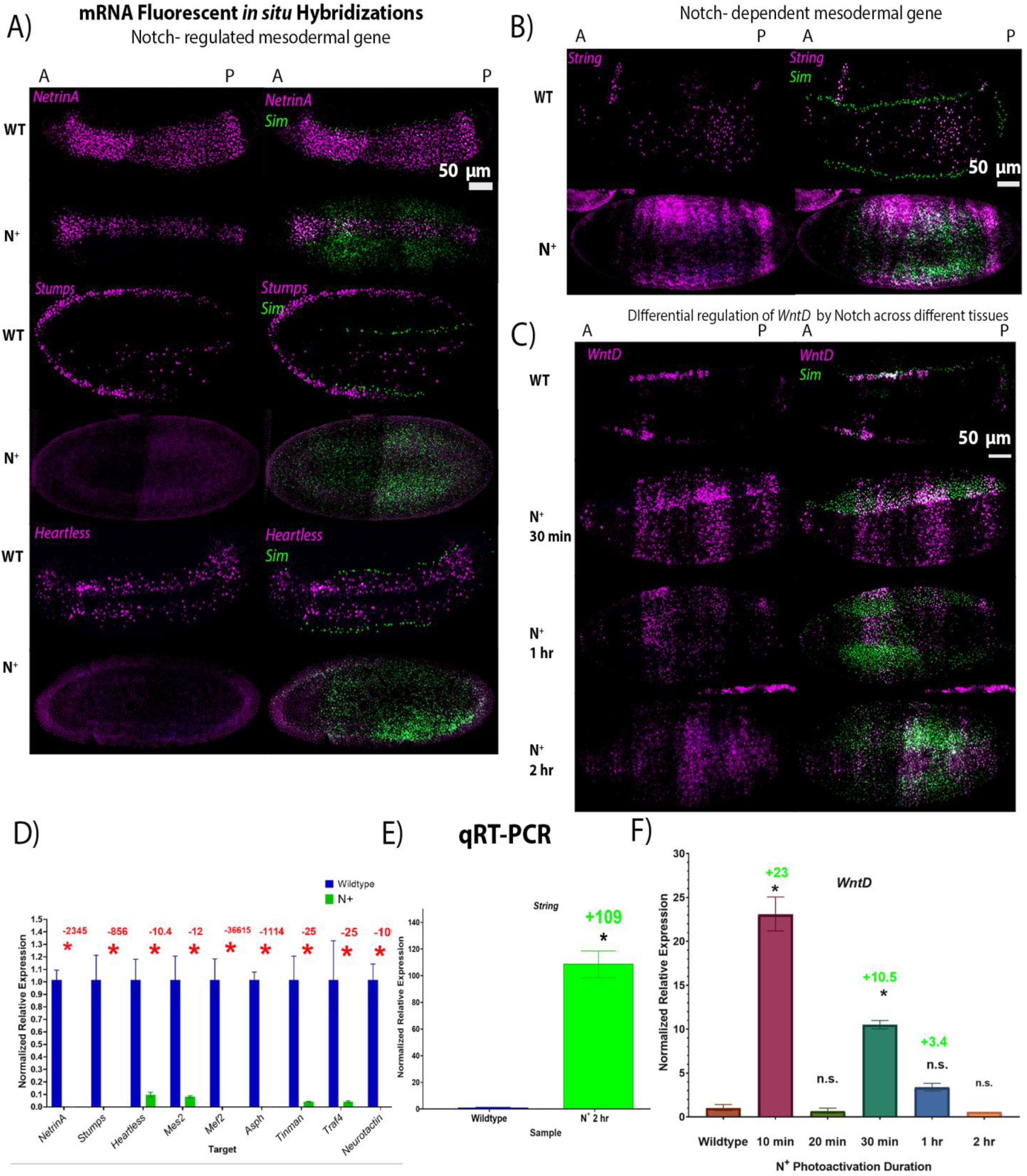
Expression of mesodermal genes in wild-type and gain-of-function Notch mutant (Opto-Notch) embryos shows that Notch suppresses most mesodermal genes and the sufficiency of Notch to drive overexpression of *WntD* and *String*. A-C) fluorescent mRNA *in situ* hybridizations against *sim* (green) and mesodermal genes (purple) in wild-type (WT) and N+ embryos (OptoNotch embryos photoactivated for 2 hours) at cephalic furrow (stage 6). A) shows examples of genes whose expression was significantly reduced in N^+^ embryos compared to their WT counterparts; *Stumps, NetrinA*, and *Heartless*. B&C) shows the expression of the two genes whose expression was upregulated in Delta mutants: *String* and *WntD*. Their expression expands laterally into the ectoderm of N+ embryos. D-F) qRT-PCR generated normalized relative expression of mesodermal genes in wild-type and N^+^ embryos at cephalic furrow (stage 6). D) Expression of *Asph, Mef2, Mes2, Neurotactin, NetrinA, Stumps, Tinman, Heartless*, and *Traf4* is significantly reduced in N^+^ compared to stage-matched wild-type embryos. p-values from t-test (described below) are: *Asph*: p=1.54988×10^-06^, *Mef2:* p=5.32916×10^-07^, *Mes2:*p=0.0002, *Neurotactin:* p*=*8.46069×10^-06^, *NetrinA*: p=2.1557×10^-05^, *Stumps:*p=3.29648×10^-06^, *Tinman*: p=7.16119×10^-05^, *Heartless:* p=0.0006, and *Traf4*: p=0.0006. E) Expression of *String* is significantly increased in N^+^ embryos compared to wild-type embryos, p= 0.00028947. For E & D independent two-tailed T-tests were performed for each gene to compare expression in N^+^ to that in wild-type embryos. ‘*’ indicates statistical significance determined by a p-value < 0.05. Error bars represent the standard error of the mean. F) Expression of *WntD* is significantly increased to different levels in OptoNotch embryos photoactivated for different durations of time. N=30 embryos for each group. A one-way ANOVA was performed, p=1.63×10^-06^, F= 31.6664. A post-hoc Tukey HSD test was used to perform pairwise comparisons. Only significant differences from WT are shown here, denoted by ‘*’, which represents a p<0.05. Error bars represent the standard error of the mean.

Class 1- Genes whose expression is repressed by Notch signalling overactivation. These include *Asph, Mef2, Mes2, Neurotactin, String, Stumps, Tinman, Traf4, Twist*, and *NetrinA* (Figure 4A & D). Fold-changes are denoted in Figure 7D and p-values are reported in the figure caption.

Class 2- Genes whose expression is significantly upregulated and expanded outside of the mesoderm and mesectoderm: *String* and *WntD* (Figure 4B,C,E&F). Fold-changes are denoted in Figure 7D and p-values are reported in the figure caption.

In wild-type embryos, *WntD* is expressed at a high level in all mesectodermal cells and at a lower level in a heterogeneous segmented manner across the anterior-posterior axis of the ventral mesoderm, whereas it is not expressed in the ectoderm (Figure 4C)^[25]^. This unique expression pattern is a distinguishing feature of the mesoderm cells. In contrast, we observed that overactivation of Notch signalling in OptoNotch embryos alters *WntD* expression such that the mesectodermal cells exhibit lower expression levels, similar to those observed in neighbouring mesodermal cells. Strikingly, we also observed significant, titratable expansion of *WntD* expression in ectodermal cells. *WntD* expression was increased in Opto-Notch embryos that were photoactivated for 30 mins in the mesoderm but was still segmented and this pattern was distinct from that of neighbouring mesectoderm cells, where *WntD* was expressed in a continuous (non-segmented) 2-3 cell-wide stripe along the AP axis. In contrast, *WntD* was expressed in mesodermal, mesectodermal, and ectodermal cells in a segmented pattern across the AP axis of Opto-Notch embryos photoactivated for 1 hour, which exhibited the highest levels of Notch signalling, as measured by increases in *sim* expression (Figure 3). Ectopic Notch signalling alters the expression profile of mesodermal, mesectodermal and ectodermal cells and therefore these cells are less distinguishable from one another. This emphasizes the role of Notch signalling in boundary refinement.

Our observation that mesoderm and mesectodermal/ectodermal expression of *WntD* was differentially regulated by Notch signalling drove us to further investigate the regulatory relationship between Notch signalling and *WntD* expression levels. Given that *WntD* is a transcriptional target of the Dorsal/Twist/Snail network, we decided to investigate whether changes in the levels of Notch signalling activation affected the expression of Twist and Snail^[25]^.

### 2.5. Notch signalling regulates *Twist* and *Snail* expression

Twist and Snail are both transcription factors that play a critical role in mesoderm patterning and in gastrulation^[2,4,26]^. Twist is a transcription factor that is necessary for the expression of most mesodermal genes involved in gastrulation, while Snail is a transcriptional repressor that inhibits the expression of genes that specify other tissues, such as the mesectoderm and the ectoderm^[4,26]^. We analyzed embryos at cellularization to determine the number of cells in the mesoderm, cells expressing Twist.

We found that the number of Twist-positive cells of loss-of-function Delta mutants (Dl^-^) and gain-of-function (N^+^) Notch mutants is significantly reduced from 18 cells (± 0.8, n=5) in wild-type to 12 (±0.86, n=4) in Dl^-^ and 9 (± 0.67, n=3) in N^+^ mutants (Figure 5A, 5B). Ventral furrow invagination proceeds normally in Dl^-^ mutants, but in N^+^ mutants, there are additional groups of cells that invaginate, giving rise to irregular, incomplete invagination in the embryos and the ventral furrow appears disorganized following ingression (Figure 5A, D). During later stages of gastrulation, both Dl^-^ and N^+^ mutants show developmental defects, which resemble the Twist loss-of-function mutant phenotype (Figure S1). The reduced protein expression of Twist and Snail is supported by the reduced gene expression of these mutants compared to their wild-type counterparts (Figure 5C).

**Figure 5.**
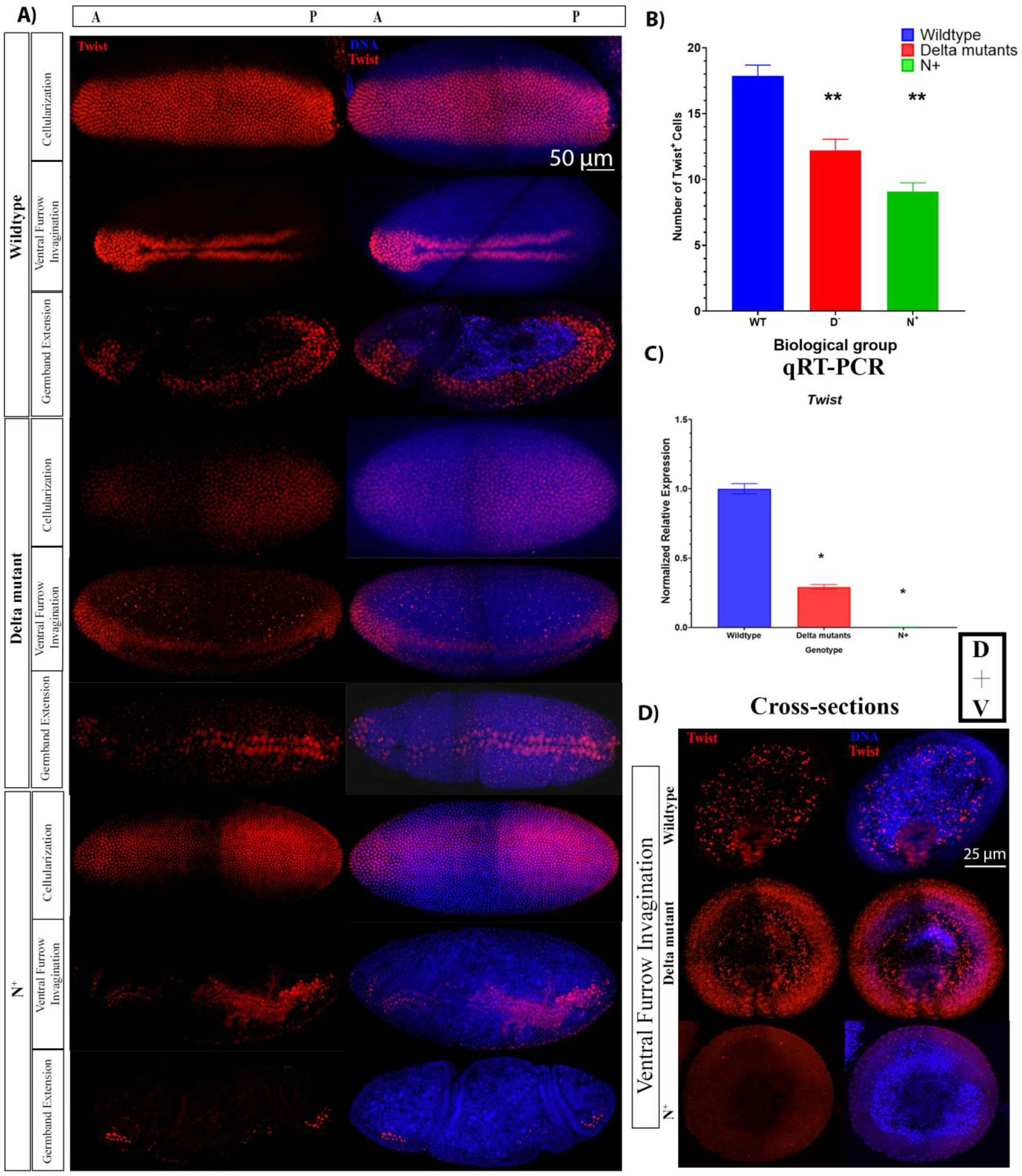
Twist protein and mRNA expression varies between wild-type and Notch gain-of-function and loss-of-function mutant embryos. A) Twist protein expression during cellularization, early and late gastrulation in wild-type (WT), loss-of-function (Delta mutant (Dl^-^)) and gain-of-function Notch mutants (N^+^, Opto-Notch embryos photoactivated for 2 hours prior to fixation). B) Number of Twist positive cells in cellular blastoderms (WT: n= 4, Dl^-^: n=5, N^+^: n=3). Error bars represent standard deviation. A one-way ANOVA was used to test for significance, p<0.0001, F=126, then Tukey HSD test was performed, which showed that all groups were significantly different from one another, p=0.001 for all pair-wise comparisons. ‘**’ denotes that experimental group was different from WT and other experimental group. C) qRT-PCR generated normalized relative mRNA expression of *twist*, n= 30 embryos at cephalic furrow formation in each genotype. Error bars represent the standard error of the mean. A one-way ANOVA was used to test for significance, p=3.33×10^-09^, F=436.7164, then Tukey HSD test was performed, which showed that all groups were significantly different from one another, p=0.001 for all pair-wise comparisons. ‘*’ denotes that experimental group was different from WT and other experimental group. D) Twist protein expression in cross-sections of wild-type, Delta mutant and N^+^ embryos undergoing ventral furrow invagination. Twist is expressed in the invaginating cells of Wildtype and Delta mutants, however in N+ embryos, expression is reduced, and there are multiple groups of cells invaginating or beginning to invaginate.

We also analyzed Snail expression at the protein and mRNA levels. The number of Snail-positive cells during cellularization was significantly reduced in Dl^-^ (12 cells ± 1.62, n=5), and N^+^ mutants, (9 cells ± 2.43, n= 3), compared to wild-type embryos, (18 cells ±0.79, n = 4) (Figure 6A&B). In Dl^-^ mutants, *snail* mRNA expression is significantly increased while in N^+^ mutants, *snail* mRNA expression is significantly decreased (Figure 6C &D). Altogether, this indicates that Notch signalling negatively regulates *snail* expression. Given that Twist and Snail are markers of mesodermal cell fate and that the number of Twist-positive cells approximately corresponds to the number of Snail-positive cells in wild-type and mutant embryos, these findings indicate that decreases and increases of Notch signalling beyond physiological levels leads to a reduction in the number of presumptive mesodermal cells in blastoderm embryos, which leads to more drastic defects of gastrulation in older embryos (Figure 5A & 6A, Figure S1). The discrepancy between Snail protein and mRNA expression in Dl^-^ mutants indicates that there are two distinct inputs from Notch signalling, which will be discussed further below. Taken together, these data support the hypothesis that Notch signalling is active in the ventral mesoderm and is required for delineating the pattern of mesodermal gene expression and proper progression through gastrulation.

**Figure 6.**
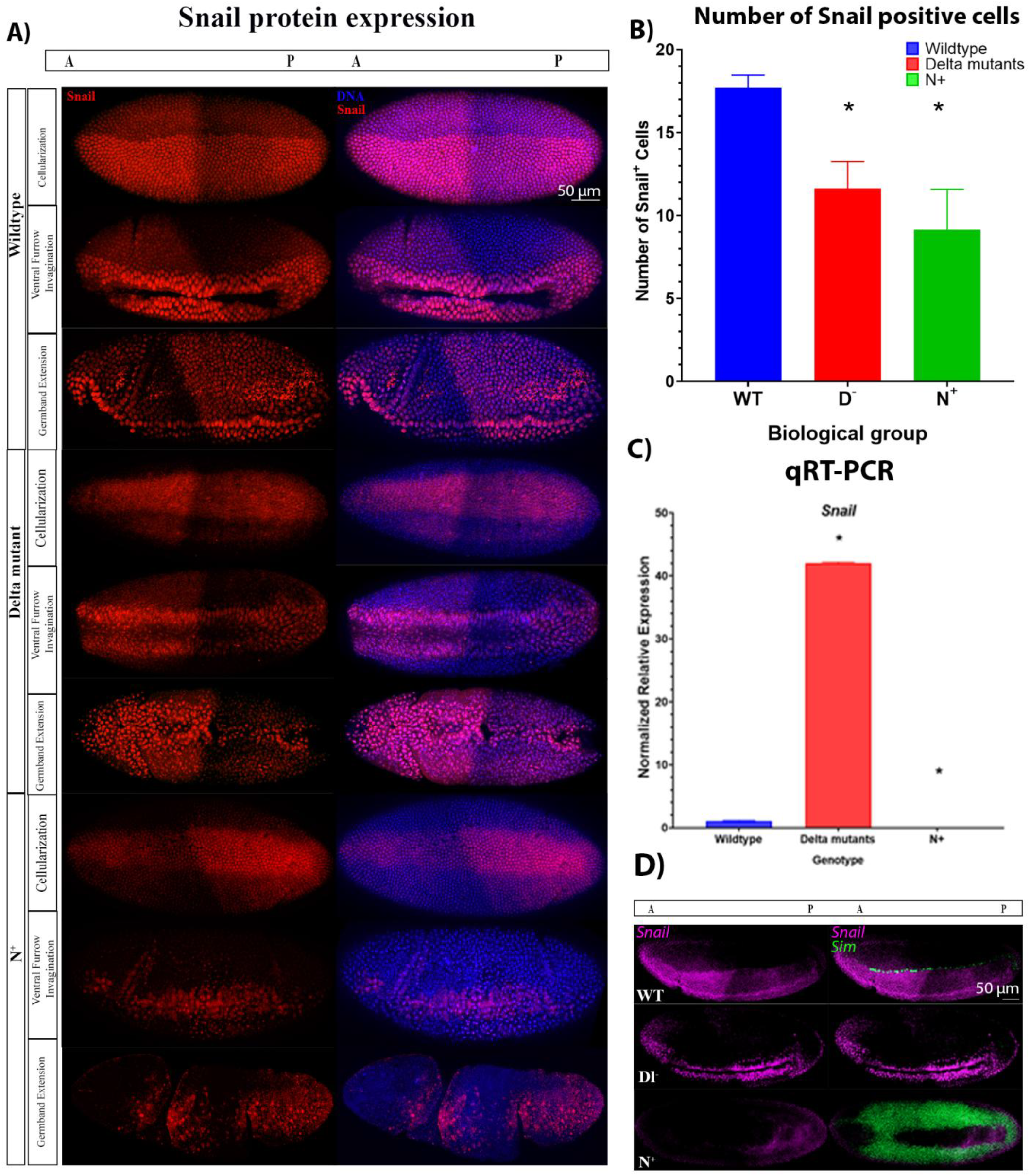
Snail protein and mRNA expression varies between wild-type and Notch gain-of-function and loss-of-function mutant embryos. A) Snail protein expression during cellularization, early and late gastrulation in wild-type, Dl^-^, and N^+^ mutant embryos. B) Number of Snail-positive cells in cellular blastoderms (WT: n= 6, Dl^-^: n=5, N^+^: n=3). Error bars represent standard deviation. One-way ANOVA was used to determine whether there were significant differences, p=0.0002, F= 26.24, then Tukey HSD test was performed, which showed that Dl^-^ (p=0.001) and N^+^ (p=0.001) were significantly different from WT embryos but not from one another (p=0.1504). C) Normalized relative mRNA expression of *snail*, n= 30 embryos at cephalic furrow formation in each genotype. Error bars represent SEM. Statistical significance is denoted by ‘*’ determined by ANOVA (p=3.13×10^-09^) and a Tukey HSD test, which showed that all groups were significantly different from one another (p=0.001 for all pair-wise comparisons). D) mRNA fluorescent *in situ* hybridizations against *Snail* (purple) and *sim* (green) in wild-type, Delta mutant and N^+^ embryos.

Given that Twist and Snail are transcriptional targets of Dorsal, these results motivated us to investigate whether there is any input from Notch signalling on the Dorsal patterning gradient. We then analyzed the expression of Dorsal activity during cellularization in wild-type, Notch loss-of-function, and Notch gain-of-function embryos.

### 2.6. Notch signalling provides a negative feedback signal on Dorsal distribution by regulating *WntD* expression

As mentioned previously, Dorsal is a maternally deposited transcription factor that forms a morphogen gradient across the dorsal-ventral axis and is critical for mesoderm specification. In the absence of Toll signalling, Dorsal is bound to a protein named Cactus in the cytoplasm, which is responsible for restricting its entry into the nucleus^[27]^. Once Toll signalling is activated by Spätzle (Spz), Cactus is targeted for destruction, liberating Dorsal which then translocates to the nucleus, where it activates gene expression^[9,10]^.

As previously reported, we observed that the nuclear abundance of Dorsal is most concentrated in the ventral-most region of the blastoderm in wild-type embryos, which has been shown to activate transcription of two ‘master regulators’ of gastrulation: Twist and Snail^[2,4,26,28]^. In contrast, we observed that the number of cells that exhibit prominent nuclear localization of Dorsal is significantly reduced from (17 cells ±2.82, n=4) in wildtype embryos to (13 cells ± 1.46, n=3) in Dl^-^ mutants (Figure 7A, 7B). Interestingly, we observed two distinct classes of N+ mutant embryos:

1. ‘N^+^ high’, which contained more cells with a high nuclear concentration of Dorsal, 10 cells (+ 1.7, n=3) (Figure 7A&B, ‘N^+^ high’) and,
2. ‘N^+^ low’, which contained fewer cells with a high nuclear concentration of Dorsal, 3 cells (+0.48, n=3), and therefore, a narrower mesoderm (Figure 7A&B, ‘N^+^ low’).

**Figure 7.**
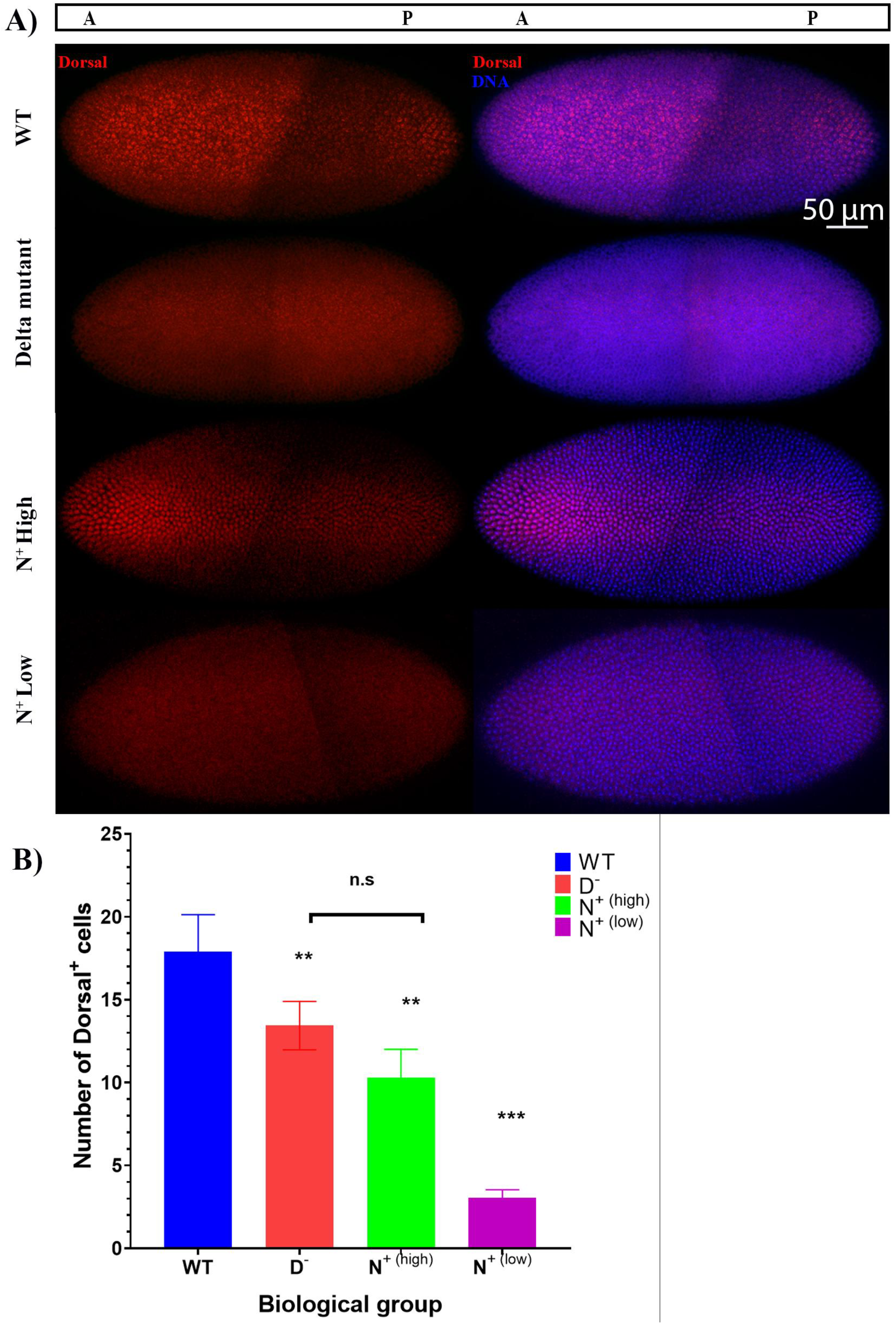
Nuclear concentration of Dorsal on the ventral side of embryos decrease in Notch gain-of-function and loss-of-function mutants compared to wild-type embryos. A) Dorsal protein expression (red) in cellular blastoderm embryos. N^+^ are Opto-Notch embryos photoactivated for 2 hours prior to fixation. B) The number of cells in the mesoderm determined using the peak region from the line scan plots. One-way ANOVA was used to test for statistical significance between wild-type (WT) embryos (n=4), Delta mutants (Dl^-^) (n=3), N^+(high)^ mutants that had a higher level of Dorsal expression (n=3), and N^+(low)^ mutants with a very low level of Dorsal signal (n=3), p<0.0001 and F=45.96. Tukey HSD test was used to perform pairwise comparisons, which showed that Dl^-^ and N^+(high)^ were significantly different from WT and from N^+(low)^, denoted by ‘**”, but not from one another, shown by ‘n.s’. N^+(low)^ was significantly different from all other groups, denoted by ‘***’. WT vs Dl^-^ p=0.0365, WT vs N^+(high)^ p=0.0017, WT vs N^+(low)^ p=0.001, Dl-vs N+(high) p= 0.1545, Dl^-^ vs N^+(low)^, p= 0.0010053, N^+(high)^ vs N^+(low)^ p= 0.0024. Error bars represent standard deviation.

This suggests that Notch signalling negatively regulates Dorsal distribution in the mesoderm as when Notch signalling is increased, fewer cells contain a high nuclear concentration of Dorsal protein. These results are consistent with our observation that the reduced expression of both Twist and Snail protein levels in Notch loss-of-function and gain-of-function mutants (Figure 5&6) as they are consistent with the fact that the decrease in the nuclear accumulation and activity of Dorsal in the ventral mesoderm results in reduced expression of its target genes, *Snail* and *Twist*. Considering that Dorsal activity is thought to precede the onset of Notch signalling during early embryogenesis, we propose that Notch is acting on the Dorsal/Twist/Snail network through an intermediate negative regulator, specifically WntD. Here we provide evidence that Notch signalling upregulates *WntD* expression. As WntD inhibits Dorsal nuclear localization, the increase of *WntD* expression leads to decreases in Dorsal nuclear localization and transcriptional activation of target genes including *Twist* and *Snail* ^[25]^. We, therefore, propose that ectopic Notch signalling is sufficient to alter ventral mesodermal patterning through the effects of it’s downstream target genes (i.e., *WntD*) on the Dorsal/Snail/Twist patterning network. Similarly, our findings indicate that decreases in Notch signalling result in reductions of Dorsal, Twist, and Snail protein expression, leading to a reduction in mesodermal cell number in the early embryo, and also result in a disorganized ventral furrow later in gastrulation, underscoring the necessity of Notch signalling for proper mesoderm formation and gastrulation.

## 3. Discussion

Disentangling the role of Notch signalling during early embryogenesis is critical to understanding the mechanisms involved in tissue formation, boundary establishment and refinement. The processes involved in tissue and boundary formation are conserved in later stages of the organism’s life as in stem cell renewal and differentiation. Unravelling the interactions between different signalling pathways, such as Notch signalling and Toll signalling, is also important for understanding disease pathogenesis, where the same processes are dysregulated and lead to disease^[29]^.

The critical role of Notch signalling during embryogenesis has long been appreciated in the context of neurogenesis and myogenesis, which both occur after early mesoderm development^[15,30]^. However, while there is evidence to suggest that Notch signalling is active in the ventral mesoderm, its role in this early embryonic tissue has been largely unexplored. Previously, there were only a few known Notch target genes that are expressed in the mesoderm, such as *Twist, String*, and *Asph*. While Notch signalling has been previously shown to regulate the expression of *Twi, Asph*, and *String* during later stages of embryogenesis and larval stages and in adult tissues, it remained unknown whether Notch signalling is required for mesodermal gene expression or mesoderm specification during early embryogenesis^[31,32]^. Here, for the first time, we present evidence for Notch activity in the ventral mesoderm and identify several target genes that are critical to mesoderm formation and gastrulation and are dependent on Notch signalling for their expression. These ventral mesoderm target genes include *tinman, Mes2*, and *Mef2*, which all encode transcription factors that are necessary for myogenesis during embryonic and larval stages^[33–38]^. Some of these target genes, whose expression was dependent at least in part on Notch signalling, include factors that are involved in orchestrating or facilitating cell shape changes that are necessary for mesoderm invagination and internalization^[39]^. These include *Traf4*, which encodes a cytoplasmic protein involved in TNF signalling that plays a critical role in cellular apical constriction and is necessary for ventral furrow invagination^[39]^. Another affected gene, *Neurotactin*, which encodes a cell adhesion molecule^[40]^. Decreases and increases in Notch signalling also affect the expression of *htl* and *stumps*, which encode key factors involved in Fibroblast Growth Factor (FGF) signalling that is required for mesoderm spreading^[11]^. Here we show that Notch signalling is required for the expression of *Stumps* but negatively regulates *htl* expression. Surprisingly, however, loss-of-function Notch mutants did not show defects in mesoderm spreading when cross-sections were examined (Figure S4). This indicates that the expression of these genes is only partially dependent on Notch signalling, which is consistent with previous studies that characterized them as targets of Twist^[36,41,42]^. Considering that Twist protein is expressed in these loss-of-function Delta mutants, we conclude that the reductions in Twist protein that we observed are not detrimental and are sufficient to support the role of Twist in regulating its downstream target genes in promoting mesoderm spreading.

To test the sufficiency of Notch signalling to drive the expression of mesodermal genes, we developed a novel light-gated tool that allows precise temporal control. This system provides several advantages compared to traditional overexpression models that have been used previously: 1) OptoNotch is titratable as shown by the incremental expansion of *sim* expression (Figure 3C&D), 2) the combination of OptoNotch with tissue-specific Gal4 drivers can provide more precise tissue-specific and temporal control, and 3) OptoNotch has the potential to provide more precise spatial control if combined with light sheet microscopy or 2-photon microscopy. In this study, OptoNotch provided titratable temporal control over Notch activation. There are a few cells in the embryo that did not express *sim* regardless of photoactivation time, which indicates that there are additional regulatory mechanisms that repress *sim* expression directly or act indirectly by inhibiting NICD translocation to the nucleus. In addition, we observed a decrease in ectopic expression after longer periods of continuous photoactivation, i.e., embryos photoactivated continuously for one hour had higher total *sim* expression than those photoactivated continuously for 2 hours and more cells, in the mesoderm and ectoderm, ectopically expressed *sim* in the former than the latter. This finding supports another report, which found that ectopic *sim* expression decreases after continuous overactivation of NICD, potentially due to nuclear desensitization to NICD^[43]^.

In this study, we present evidence that critical mesodermal genes require Notch signalling for normal expression, shown by significant decreases in mesodermal gene expression in both loss-of-function and gain-of-function Notch mutant embryos. In the gain-of-function Opto-Notch mutants that were photoactivated for a longer duration, expression of most of the mesodermal target genes was significantly reduced (Figure 4), however expression in those photoactivated for shorter durations was significantly increased for many of the candidate genes including *Mes2, Tinman, Stumps, Twist*, and *Mef2* (Figure S3). This indicates that Notch is sufficient to increase the expression of these target genes and that different genes require different levels of Notch signalling. The decrease in expression we observed in these target genes following a long duration of photoactivation is likely caused by two mechanisms: 1) negative feedback inhibition inputs that follow increases in gene expression, some of which are Notch target genes encoding gene repressors, (*E(spl)C* and potentially *emc*)^[31]^ and 2) decreases in *Twist* expression, as most of these genes are also transcriptional targets of Twist^[36]^. Our findings are supported by previous experiments performed later in gastrulation that showed that Notch signalling regulates mesodermal gene expression directly (i.e., *Twist, Asph*, and *String*) and indirectly by activating the expression of repressors of Twist^[31]^.

Some of the candidate mesodermal Notch target genes we identified were down-regulated following Notch overactivation regardless of photoactivation duration including *Asph, Neurotactin, Traf4*, and *Heartless*. Although these genes contain consensus Su(H) binding sites in their extended gene regions, our expression analysis indicates that Notch signalling is not sufficient to increase their expression in the mesoderm. This is likely due to their repression by other Notch target genes such as the *E(spl)C* genes, including *emc*, although this was not tested here. Another mechanism may involve negative feedback through post-translational modification regulation of endogenous NICD and/or OptoNotch proteins that promote their degradation^[44,45]^. In addition, post-translational modifications may also inhibit translation of particular transcripts or increase the rate of translation of others, which may explain some of the discrepancies observed between the level of RNA and protein (i.e., *snail* mRNA is increased in loss-of-function mutants but protein expression is decreased). A potential role of post-transcriptional and post-translational regulation of transcriptional activation is supported by the decrease in the fold-change of *sim* expression in embryos that are photoactivated for 2 hours compared to 1 hour (Figure 3). Taken together, our results provide clear evidence that Notch signalling is normally active in the mesoderm during early embryogenesis and is required for normal levels of mesodermal gene expression. This requirement of Notch signalling is supported by our observation of defects the ventral furrow during later stages of gastrulation (following the internalization of the mesoderm) in loss-of-function mutants (Figure 5). The ventral furrow of loss-of-function mutants is internalized normally and cross-sections show that mesoderm spreading appears normal. However, in some older mutant embryos the ventral furrow appears disorganized (it is jagged, rather than straight, and has some gaps, where the ventral closure is not complete) (Figure 5A & 6A, ‘germband extension Dl^-^ embryos). These observations indicate that the changes in mesodermal gene expression while not catastrophic for initial mesoderm formation and internalization, lead to incomplete furrow zipping/closure which then contributes to the breakdown of the rest of gastrulation.

Interestingly, we identified one gene, *string*, that behaves as a classic direct target of Notch signalling; its expression is significantly reduced in the absence of Notch signalling and is significantly increased upon overactivation of Notch signalling. *String* encodes a phosphatase, cdc25/String, that activates a mitotic kinase, cdk1^[45,46]^. Zygotic cell division is dependent on *string* expression^[45,46]^. Previous work showed that loss-of-function *string* mutants exhibit cell cycle arrest after nuclear cycle 14, which is the last cycle controlled by maternally deposited String, however, cell shape changes and morphogenetic movements are not affected except that cell numbers in the mesoderm are reduced^[45,46]^. The precise temporal expression of *string* has been shown to be critical for the coordination of cell division^[46,47]^. The morphological defects observed in the N^+^ mutants described here may also be due to premature accumulation of *string*, causing premature cell division prior to mesoderm internalization.

Interestingly, we identified a gene, *WntD*, that is differentially regulated by Notch signalling across neighbouring tissues, the mesoderm and mesectoderm. Our expression analysis of loss-of-function mutants indicates that Notch signalling is required for *WntD* expression in mesectoderm cells that flank mesoderm cells on both sides, while it negatively regulates its expression in the mesoderm, as seen by the increase in expression in this region in the absence of Notch signalling activity (Figure 2). The presence of Su(H) consensus binding sites in the *WntD* genomic region indicates that Notch can activate its expression directly by binding to Su(H). This is likely the main input on *WntD* in the mesectoderm. Our results imply that Notch activates the expression of a repressor of *WntD* in the mesoderm and in the absence of Notch signalling, this repression is relieved. The identity of this repressor remains unclear. In the gain-of-function mutant expression of *WntD* is expanded laterally into the ectoderm, much *sim* expression in those embryos and the boundary between the mesoderm and ectoderm becomes much less defined. This finding highlights the role of Notch signalling in defining the boundary between different tissues. Increases or decreases in Notch signalling, result in changes in the expression profile of these boundary cells. In the loss-of-function mutants, the boundary cells lose expression of *WntD* and *sim*, while in GOF mutants, these cells are no longer distinct from neighbouring cells, as they express these two genes at similar levels to the neighbouring mesoderm and ectoderm cells. These changes in expression profile most likely contribute to the defects in the embryonic midline observed in the GOF mutants.

*WntD* encodes a secreted ligand of the Wnt family of proteins and has been identified as an inhibitor of Dorsal^[25]^. It has been shown that *WntD* expression is regulated by the Dorsal/Twist/Snail genetic network and provides negative feedback inhibition by competitively binding to the Toll receptor instead of Spz thereby preventing nuclear transport of Dorsal^[48]^. Our findings show that increases in *WntD* expression coincide with decreases in the size of the region with the most concentrated nuclear Dorsal, supporting the mechanism of negative feedback inhibition shown previously^[48]^. As expected, the decrease in nuclear Dorsal concentration results in decreases in the number of Snail- and Twist-positive cells, indicating that the mesoderm region is narrowed in both loss- and gain-of-function mutants, which emphasizes the role of Notch signalling in the development of the mesoderm. The discrepancy between Snail protein and mRNA expression levels, being that *snail* mRNA is increased but Snail protein levels are decreased in Dl^-^ mutants indicates that there are two distinct inputs from Notch signalling on Snail expression. Notch may directly downregulate *snail* expression but simultaneously upregulates a positive regulator of Snail protein expression, either on the transcriptional or post-transcriptional level, or vice versa. Put together, these findings provide evidence that in addition to functioning in a cell-autonomous manner to drive mesodermal gene expression, Notch signalling also provides feedback inhibition signals on the upstream patterning programme, regulated by Toll signalling through Dorsal, indirectly by regulating *WntD* expression.

## 4. Materials and Methods

### Analysis of ChIP-seq data for Su(H) binding sites in mesodermal genes

We generated a list of genes that were expressed in the mesoderm and were differentially expressed at nuclear cycle 14 of embryogenesis based on mRNA expression profiles available on BDGP and RNA-seq data from the modEncode project^[23]^.

To determine Su(H) binding sites in the extended regions of the genes of interest, we analyzed ChIP-seq data generated by Ozdemir et al. (2014)^[17]^. The study generated Su(H) ChIP-seq data for 2–4-hour yw embryos using goat and rabbit antibodies. To predict Su(H) binding sites, we separately analyzed the data generated by the replicate experiments with the two antibodies and late combined the findings to get the non-redundant list of Su(H) binding sites for every gene.

The wig files were retrieved from Gene Expression Omnibus (GEO) (GEO accession: GSE59726). The wig files were converted to bigwig format which were then converted to BEDgraph format using the tools wigToBigWig and bigWigToBedGraph respectively. We further used the MACS3 peak caller to call peaks using BEDgraph files^[49]^. The 20bp region surrounding the point source of the peak was taken as the binding site. The predicted Su(H) binding sites were intersected with the extended regions of genes using BEDtools to get Su(H) binding sites on the genes’ extended regions^[50]^.

### Fly crosses and transgenic lines

Flies were maintained at 25 °C and fed with semi-defined food made as described^[51]^. Most of the stocks used here were obtained from the Bloomington Stock Center or were gifted by other labs as outlined in Table S1.

W1118 flies were used as wild-type controls. Delta-TS are heterozygous for two temperature-sensitive alleles of Delta, which are fully functional at room temperature (∼21-23 °C) but non-functional at 32 °C (Figure S5) ^[52]^. Resultant embryos from Delta-TS flies were collected for 1.5 hours, then heat-shocked at 32 °C for 2 hours and 20 minutes and then fixed and stored in methanol at -20 °C.

We used the Best Gene injection services to generate our transgenic fly lines using Φ C31-mediated attp/attB transgenesis. The mCD8-CIBN-NTEV-NICD-GFP transgene was inserted using the attP40 insertion site on chromosome 2 and balanced over CyO, while the Cry2-CTEV-mCherry transgene was inserted using the attP2 insertion site on chromosome 3 and balanced over TM3.

To generate OptoNICD embryos, homozygous UASp> Cry2-CTEV-mCherry virgin females were crossed to homozygous UASp>mCD8-CIBN-NTEV-NICD-GFP males. Males and females from the resultant generation (heterozygous for both alleles) were crossed and their progeny was screened for double homozygote virgin females which were crossed to male Mat- alpha- tublin> Gal4 flies. The females from this progeny, whose genotype was {[y1] [w*]; UASp>mCD8-CIBN-NTEV-NICD-GFP/Mat- alpha- tublin> Gal4; UASp> Cry2-CTEV-mCherry/Mat- alpha- tublin> Gal4} were then crossed to any of their male siblings and their resultant embryos, named OptoNICD were used for experiments. These embryos were collected in the dark and fixed in the dark (photoactivation time (T=0)) or collected in the dark and photoactivated for 10, 20, 30, 60 or 120 minutes before fixation. We adjusted the collection times so the embryos would be fixed around the time they reached cephalic furrow formation or be within 30 minutes of this developmental time point. In other words, the longer the exposure time, the shorter the collection time. We also collected embryos that were for 120 minutes and then aged in the dark for another 1 hour prior to fixation to study the effects of ectopic Notch activity on gastrulation progression. N^+^ embryos were photoactivated for 120 minutes before fixation or freezing for RNA collection.

### Cloning

We adapted the previously described system to develop a light-inducible variant of NICD^[53]^. Our system is composed of two distinct proteins. One is a cytosolic protein containing Cry2, mCherry, and the C-terminal portion of a split TEV protease (Cry2-mCherry-CTEV), while the other is a membrane-tethered protein containing mCD8, CIBN, the complementary N-terminal portion of TEV protease, an ASLOV domain which masks a TEV cleavage sequence in the dark that precedes NICD, and GFP (mCD8-CIBN-NTEV-NICD-GFP). Upon illumination, Cry2 binds to CIBN, bringing the two components of the TEV protease together to reconstitute the functional protease and ASLOV undergoes a conformational change which exposes the TEV cleavage sequence. The TEV protease then cleaves its cleavage sequence liberating NICD from the membrane. NICD contains a nuclear localization sequence as the endogenous protein does, so it translocates to the nucleus once it is liberated from the membrane.

We generated the Cry2-mCherry-CTEV cassette using PCR and Gibson assembly then subcloned it into a pUASP-attb plasmid (DGRC # 1358). The resultant plasmid is named pUASP-attb-Cry2-CTEV-mCherry. The sequences were amplified from the following plasmids: pCRY2PHR-mCherryN1 (addgene# 26866) and FRB_SCF2_LD6_CTEV (addgene# 58879). The primers used were purchased from Sigma and are listed in Table S2. We also subcloned this cassette into a pAc5.1/V5-His A plasmid (Invitrogen #V4110-20) by cutting the cassette out of pUASP-attb-Cry2-CTEV-mCherry using NotI and XbaI (NEB #R3189S and #R0145S) and ligating to the linearized pAc5.1/V5-His A. The resultant plasmid was named pAC5.1-Cry2-mCherry-CTEV. Similarly, we subcloned the mCD8-CIBN-NTEV-NICD-GFP cassette, which was synthesized by Biobasic, into pUASP-attb and pAC5.1/V5-His A. The resultant plasmids were named: pUASP-attb-mcd8-CIBN-NTEV-NICD-GFP and pAC5.1-mcd8-CIBN-NTEV-NICD-GFP, respectively. We generated a pUASP-attb-mcd8-CIBN-NTEV-NICD without the GFP tag to allow us to perform experiments with other GFP-tagged proteins. We also inserted these cassettes into pUAST-attb (DGRC #1419) for future use. The resultant plasmids were named: pUAST-attb-mCD8-CIBN-NTEV-NICD-GFP and pUAST-attb-Cry2-CTEV-mCherry.

### S2 cell culture and Transient Transfection

We used S2-DRSC cells (DGRC # 181) cells to test our Optogenetic tool prior to generating transgenic fly lines. S2 cells were cultured at 25 °C in Schneider’s Insect Medium (Sigma-Aldrich Co. LLC #S0146) containing 10% heat-inactivated fetal bovine serum (FBS), 500 ng/mL of insulin, and 100 µg/mL of Penicillin and Streptomycin. To transiently express our Optogenetic variant of NICD, we transfected S2 cells with pAC5.1-mcd8-CIBN-NTEV-NICD-GFP and pAC5.1-Cry2-mCherry-CTEV using the TransIT-Insect Reagent (Mirus, MIR #6104) according to manufacturer’s recommendation. The media was changed 24 hours after transfection and cells were imaged between 36 and 48 hours after transfection. Cells were kept in the dark until imaging.

### qRT-PCR

For each genotype, 30 embryos were staged, and hand selected at cephalic furrow, prior to complete invagination of the ventral furrow. We used this developmental stage for several reasons; 1) the genes of interest are all expressed zygotically and maternal material has been degraded, 2) Notch signalling is active, 3) this stage represents early gastrulation and 4) limits variability, since ventral furrow invagination occurs over 15 mins in wild-type embryos. Embryos were flash frozen in liquid nitrogen and stored at -80 °C for no longer than 1 week before RNA extraction. Total RNA was extracted using Trizol (Molecular Research Center Inc #TR 118) according to a previously published protocol^[54]^. 1ug of RNA was used to synthesize cDNA using (Bio rad kit) according to manufacturer’s instructions. 1 ul of the synthesized cDNA was used in 10 ul qPCR reaction, assembled using SYBR green (bio rad kit) according to manufacturer’s instructions. CFX maestro (machine) was used to read samples. Reactions were run in triplicates and negative controls, no-reverse transcriptase control and no-template controls were included in each run. Gene expression in Delta-loss-of function mutants was compared to gene expression in wild-type embryos. Gene expression in OptoNICD embryos expressing both components of the OptoNotch: UASp>mCD8-CIBN-NTEV-NICD-GFP and UASp> Cry2-CTEV-mCherry were compared to embryos expressing UASp>mCD8-CIBN-NTEV-NICD-GFP to account for any changes in global expression induced by the overexpression system. Data was analyzed using the relative quantification method and normalization was performed using *RPS20* and *RPL32* as housekeeping genes. Primers used for each target gene were purchased from Sigma and are listed in Table S3. For experiments where only two genotypes were compared, significance was determined using two-tailed T-tests, performed using Excel, a p-value < 0.05 was considered significant and denoted by “*”. For experiments, where more than two genotypes were compared, a one-way ANOVA was performed to determine if differences were significant, determined by a p-value <0.05. If the p-value was <0.05, a post-hoc Tukey HSD test was performed to determine which groups were significantly different from one another.

### Immunohistochemistry

Embryos were collected on apple juice agar plates and fixed in 4% paraformaldehyde (Electron Microscopy Sciences, cat. 15710) and permeabilized in 1X PBT. Embryos were incubated to in 5% blocking buffer (5% skim milk in 1X PBT) for 20 minutes. Primary antibodies were diluted in 1% blocking buffer as follows; α-Twist (1:100) (kindly gifted by Eric Wieschaus), α-snail (1:100) (kindly gifted by Eric Wieschaus), α-Dorsal (1:20, DSHB), α-NECD (1:500, DSHB, C458.2H), and α-Delta (1:20, DSHB, C594.9B), added to the embryos and then incubated at 4 °C overnight. The next day, following three washes with 1% blocking buffer, the secondary antibodies were diluted as follows and added to the embryos, which were rocked at room temperature for 2 hours; α-mouse Alexa 568, α-mouse Alexa 488, and α-rat Alexa 568. Then the embryos were washed with 1% blocking buffer twice followed by a wash with 1X PBT, one wash with 1X PBS and then DAPI was diluted in 1X PBS and added to the embryos which were then rocked at room temperature for 20 minutes. Embryos were washed twice with 1X PBS and mounted in 50% glycerol/1X PBS. Cross-sections were cut by hand using a razor blade.

### Fluorescent mRNA in situ hybridization

Anti-sense probes were synthesized using the DIG and Biotin-RNA labelling kit (Roche, 11277073910 and 11685597910, respectively) according to manufacturer’s recommendations. DNA templates were amplified using a forward primer and reverse primer containing a T7 RNA polymerase promoter sequence (5’-GTAATACGACTCACTATA<u>G</u> -3’) from cDNA clones were obtained from DGRC (Table S4). A probe against *Sim* was labelled with Biotin and the probes against the mesodermal genes were labelled with DIG to allow us to perform dual-probe *in situ* hybridizations. *In situs* were performed according to previously described protocol with the following modifications^[55]^. The hybridization buffer was supplemented with 5% Dextran Sulfate (Sigma #D6001). Rather than using milk to block, we used 5% BSA for the initial blocking step and 1% for the washes in between antibodies and diluted the antibodies in 1% BSA. We incubated embryos in TSA buffer (Tyramide Cy3 AATBioquest Sciences 11065, in 0.003% H_2_O_2_) for 1 hour, except when probing for *wntD* and *Snail*, where a 2-hour incubation was necessary. The following antibodies were used in the following dilutions: monoclonal mouse α-biotin (1:300) (Jackson Immunoresearch, 200-472-211), Anti-Digoxigenin-POD Fab fragments (1:100) (Roche, 11207733910), and Goat α-mouse Alexa 488 (1:300) (Invitrogen A32723TR).

### Microscopy

All the imaging in this paper was performed using an inverted Zeiss Axio Observer spinning disc confocal microscope equipped with a Yokogawa spinning disc head and a Prime BSI 16-bit camera fitted with 4 laser lines (350-400 nm, 450-490 nm, 545-575 nm, 625-655nm). Samples were acquired on a 20X objective (0.8 NA) for whole-mount embryos or a 40X oil immersion objective (1.4 NA) for S2 cells and embryo cross-sections. For experiments, where more than two genotypes were compared, a one-way ANOVA was performed to determine if differences were significant, determined by a p-value <0.05. If the p-value was <0.05, a post-hoc Tukey HSD test was performed to determine which groups were significantly different from one another.

### Image analysis

Image analysis was performed using ImageJ (RRID:SCR_002798). To quantify the number of cells in the mesoderm expressing Dorsal, Snail and Twist, we used line scans across the dorsoventral axis of embryos to generate a plot profile of the intensity across the embryo. We sampled 4 regions along the anterior-posterior axis per embryo and analyzed 3 or more embryos per genotype for each protein of interest. The specific numbers are stated in the corresponding figure captions. As expected, the plot profile generated a curve, where the peak region corresponds with the mesoderm, the area with the highest expression of these proteins. We then calculated the distance of the peak region of the plot profile and divided it by the average cell size (5 microns) to determine the number of cells that were in the peak region. We automated this process using a Python script and used it to analyze the outputs generated by ImageJ from the plots. Statistical significance was determined using a one-way ANOVA that was performed to determine if differences were significant, determined by a p-value <0.05. If the p-value was <0.05, a post-hoc Tukey HSD test was performed to determine which groups were significantly different from one another.

## 5. Conclusions

In conclusion, here we show that Notch signalling plays a dual role during early mesoderm development in embryogenesis. Notch activity drives gene expression of critical mesoderm genes and simultaneously provides negative feedback regulation by activating the expression of at least one inhibitor of mesodermal gene expression, *WntD*, to maintain appropriate levels of mesodermal gene expression. In addition, we describe the generation of a novel light-gated tool, OptoNotch, that allows precise temporal activation of Notch signalling. Lastly, we present evidence of a negative feedback inhibition loop regulated by Notch signalling acting on Dorsal, as one of the key players specifying the dorsal-ventral axis and patterning of the ventral mesoderm through its genetic regulation of Twist and Snail, key regulators of gastrulation. We, thus, identified a novel role for Notch signalling in boundary refinement and mesoderm patterning.

### Future directions and limitations

As a follow-up study, it would be interesting to identify the potential repressor that is activated in the mesoderm by Notch signalling to repress the expression of *WntD*, as well as continue to characterize the morphological defects observed in the gain-of-function and loss-of-function mutant Notch embryos. The latter can be achieved by analyzing the expression of cell adhesion molecules and cytoskeletal proteins that are critical to mesoderm internalization and spreading.

## 6. Additional Sections

### Supplementary Materials

Supplemental information can be found below.

### Author Contributions

Experiments were designed by M.M and A.N. M.M and A.N wrote the original draft of the manuscript. M.M prepared all figures. G.F cloned pUASP-Cry2-CTEV-mCherry and advised on image analysis. M.M cloned the other plasmids. A.A and A.N performed the *in silico* analysis of gene expression. A.A. wrote the methods section titled “Analysis of ChIP-seq data for Su(H) binding sites in mesodermal genes”. All other experiments were carried out by M.M. Data was interpreted by M.M, G.F, A.A, and A.N. P.L provided resources for the *in silico* analysis and supervised A.A’s work and manuscript editing. M.M, G.F., A.A, R.D.H, P.L, and A.N contributed to the preparation of the final manuscript and approved revisions. This work was supervised by A.N. Funding for this work was acquired by A.N. A.A.’s bioinformatics work was performed on the high-performance computing servers provided by SHARCNET (sharcnet.ca/) and the Digital Research Alliance of Canada (alliance can.ca).

### Funding

This research was funded by the National Science and Engineering Research Council of Canada (Grant #: RGPIN-2018-06781 NSERC).

### Institutional Review Board Statement

For all experiments performed in this study, Research Ethics Board Approval was obtained from Brock University. No vertebrates were used in this study. All methods were reported in accordance with ARRIVE guidelines.

### Data Availability

All data generated in this study is available upon request by contacting the corresponding author (Aleksandar Necakov)

### Conflicts of Interest

The authors declare no conflicts of interest.

## Abbreviations

The following abbreviations are used in this manuscript:

FGF: Fibroblast Growth Factor
Dl: Delta
NICD: Notch Intracellular Domain
Su(H): Suppressor of Hairless
*sim*: *Single minded*
nc: Nuclear Cycle
FISH: Fluorescent *in situ* Hybridization
RT-qPCR: Reverse Transcriptase-qPCR
OptoNotch: Optogenetic Notch
GFP: Green Fluorescent Protein
AP: Anterior-posterior
Dl^-^: Delta loss-of-function mutant
WT: Wild-type
N^+^: Notch gain-of-function (OptoNotch)
Twi: Twist
Sna: Snail
LOF: Loss-of-function
GOF: Gain-of-function
ChIP-seq: Chromatin Immunoprecipitation sequencing

## Appendix A

### Appendix A.1

**Supplementary figure 1.**
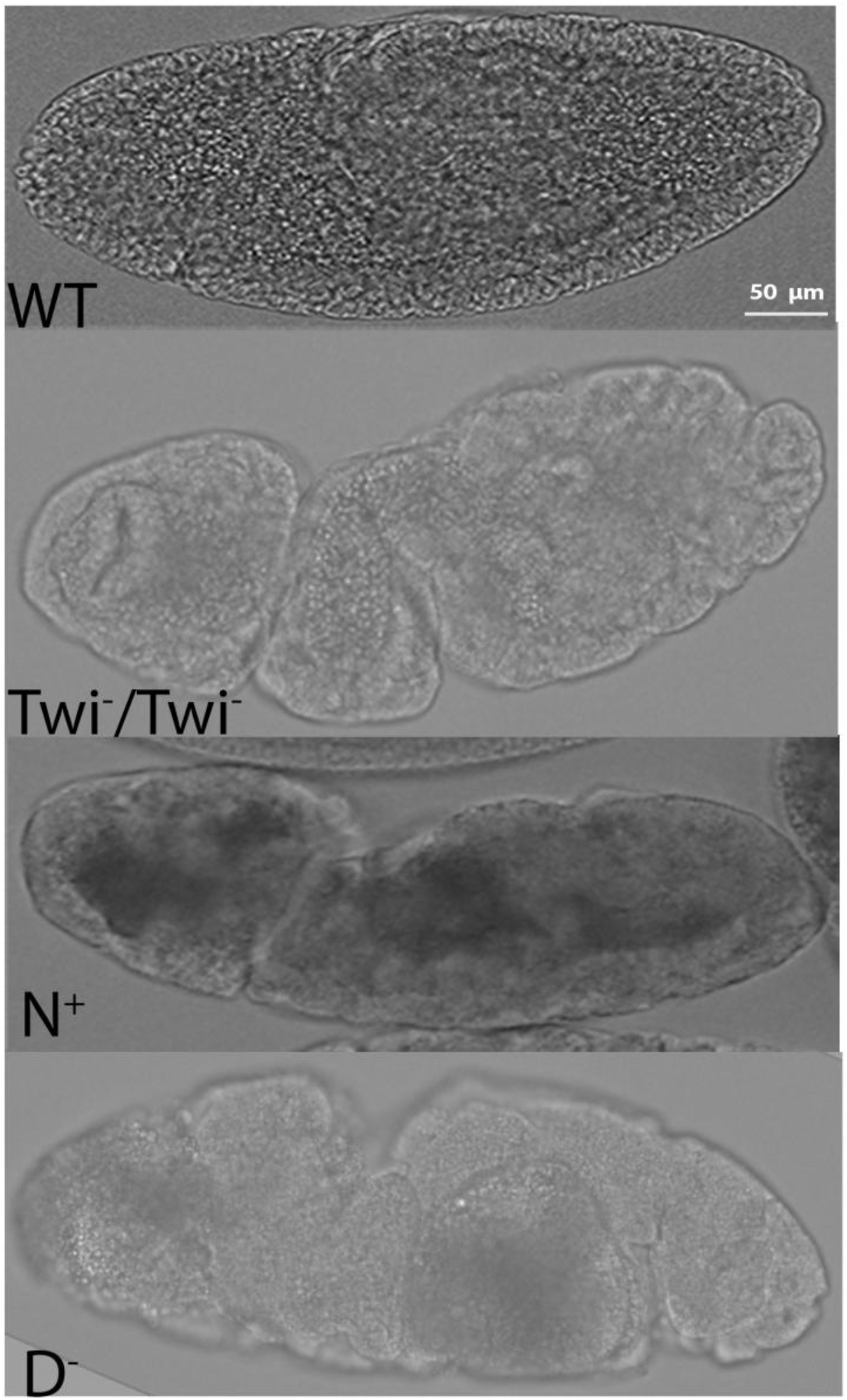
Morphological defects observed in Delta^-^ and N^+^ mutants are similar to those of Twist^-^ mutants. Bright field images of stage 9-11 wild type embryo (top), loss-of-function Twist mutant (2^nd^ image), OptoNotch mutant (3^rd^ image) and Delta^-^mutant (4^th^ image). Loss-of-function Twist mutants have similar morphological defects as gain-of-function and loss-of-function mutants in the Notch signalling pathway such as aberrant invaginations along the AP axis and the overall twisted body phenotype.

**Supplementary figure 2.**
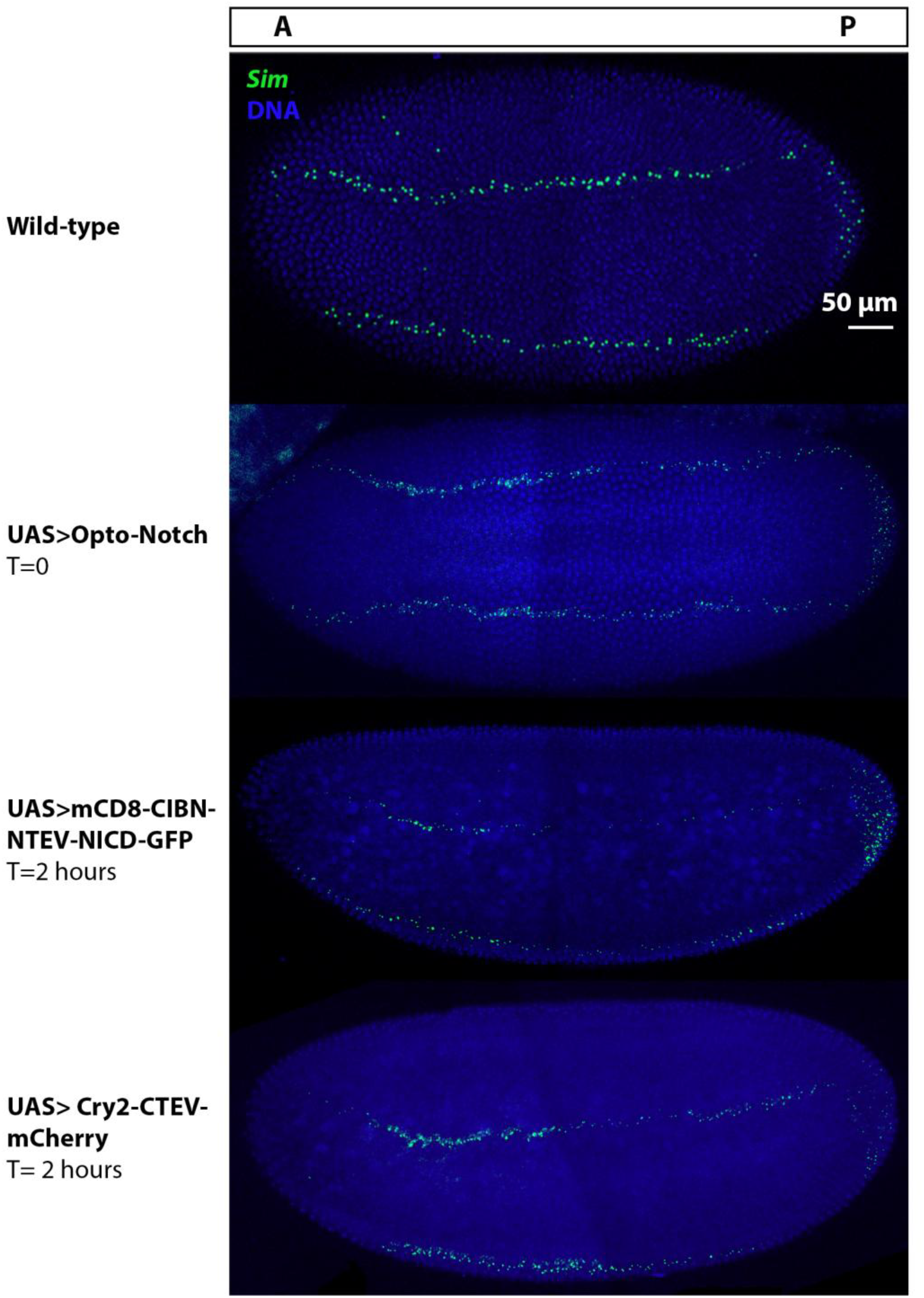
*Sim* mRNA expression does not vary between wild-type, non-photoactivated OptoNotch and control OptoNotch embryos. Fluorescent *in situ* hybridizations showing the expression of *sim* (green) in stage 5-6 embryos that; 1) do not express any transgenes (wild-type), 2) express OptoNotch but not photoactivated, 3) express only mCD8-CIBN-NTEV-NICD-GFP and photoactivated for 2 hours continuously, and 4) express only Cry2-CTEV-mCherry and photoactivated continuously for 2 hours. In embryos of all genotypes, *Sim* is expressed in the two rows of mesectoderm cells flanking the mesoderm and cells in the posterior end as in wildtype embryos.

**Supplementary figure 3.**
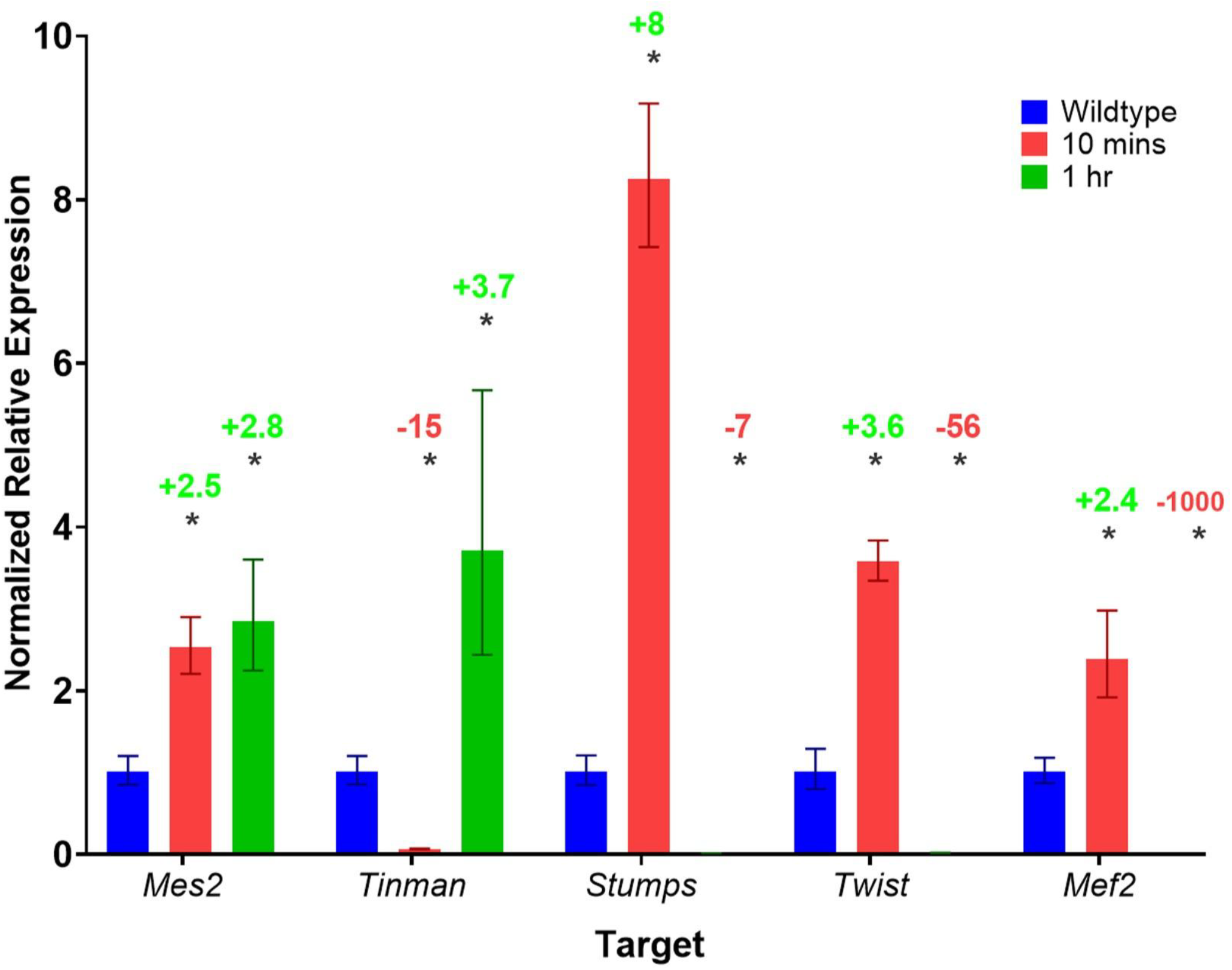
Expression of mesodermal genes in wild-type and OptoNotch embryos following photoactivation for short durations. qRT-PCR generated Normalized relative mRNA expression of mesodermal genes, n= 30 embryos at cephalic furrow formation in each genotype. Error bars represent SEM. Gene expression in wild-type (blue bars), OptoNotch embryos photoactivated for 10 minutes (red bars) and 1 hour (green bars) are shown. Short-term activation (10 mins of photoactivation) of Notch signalling results in upregulation of *Mes2, Stumps, Twist*, and *Mef2*. Long-term (1 hour of photoactivation) activation of Notch signalling results downregulation *Stumps, Twist*, and *Mef2* <u>but upregulation of</u> *<u>Mes2</u>*. Upregulation of *Tinman* expression is only observed after long-term activation of Notch signalling (after 1 hour of photoactivation). a One-way ANOVA and a Tukey HSD test. The statistical results are listed here. *Mes2*: ANOVA p=1.90×10^-08^, F=69.7468, *Tinman*: p=1.94×10^-08^, F=69.5154, *Stumps:* p=3.27×10^-11^, F=207.0422, *Twist:* p=7.80×10^-13^, F=388.3925, *Mef2*: p=6.33×10^-15^, F=870.837. Statistical significance of the difference between the experimental group and the control group (WT) is denoted by ‘*’, indicating Tukey HSD p-value <0.05 for that pairwise comparison. Only significant differences between experimental groups and WT are shown for simplicity.

**Supplementary figure 4.**
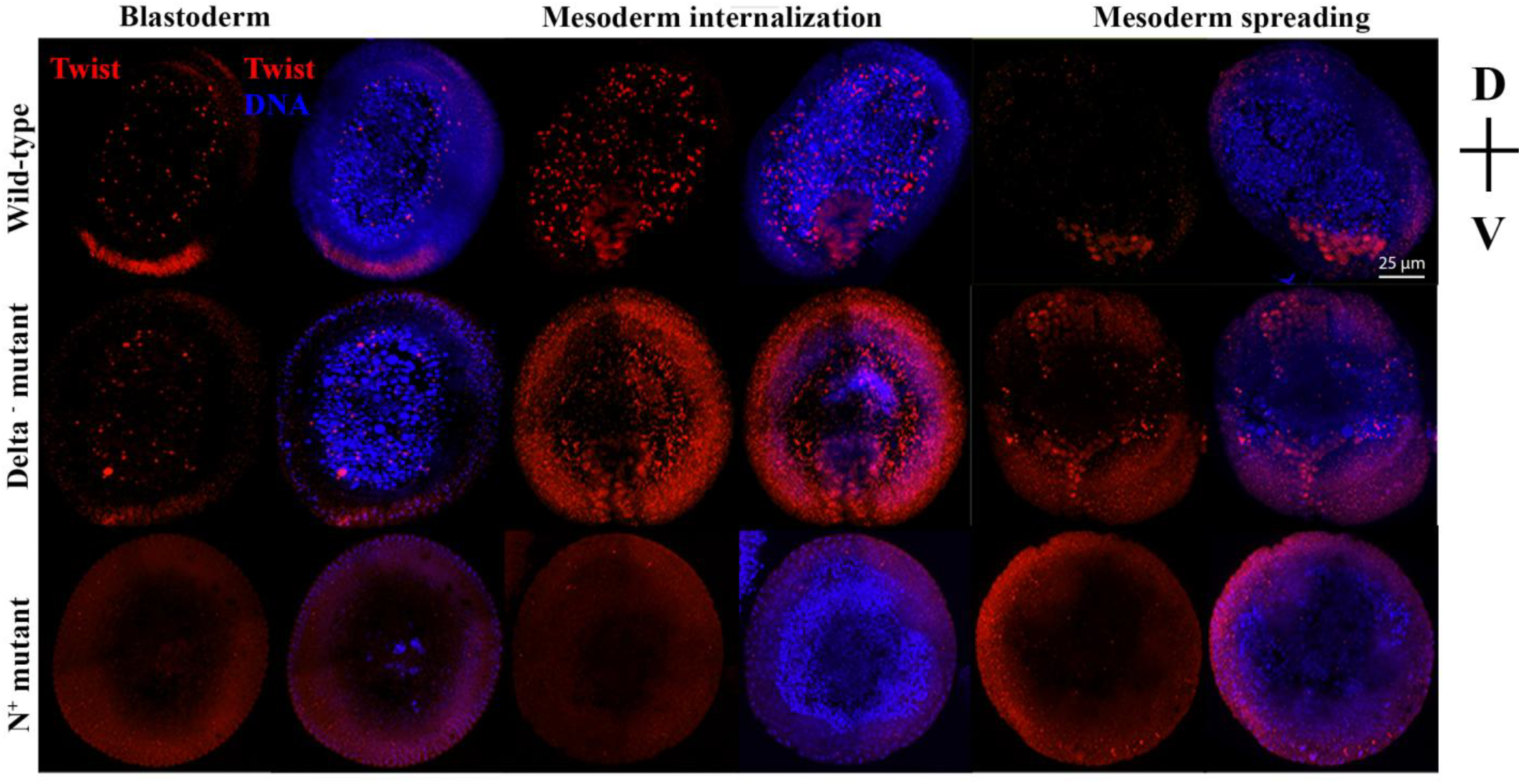
Cross-sections of embryos at different stages of gastrulation showing the decrease in Twist expression in Delta^-^ and OptoNotch (N^+^) mutants compared to wild-type embryos. Twist expression is decreased in loss-of-function and gain-of-function mutants compared to wild-type embryos during all the stages of gastrulation examined; cellularization, mesoderm internalization, and mesoderm spreading. The ventral side of embryos are at the bottom of each image. Although Twist expression is decreased, internalization and spreading of the mesoderm does not appear to be significantly impacted in mutant embryos except that in N^+^ embryos, there are additional groups of cells on the dorsal side of the embryo that are internalized.

**Supplementary figure 5.**
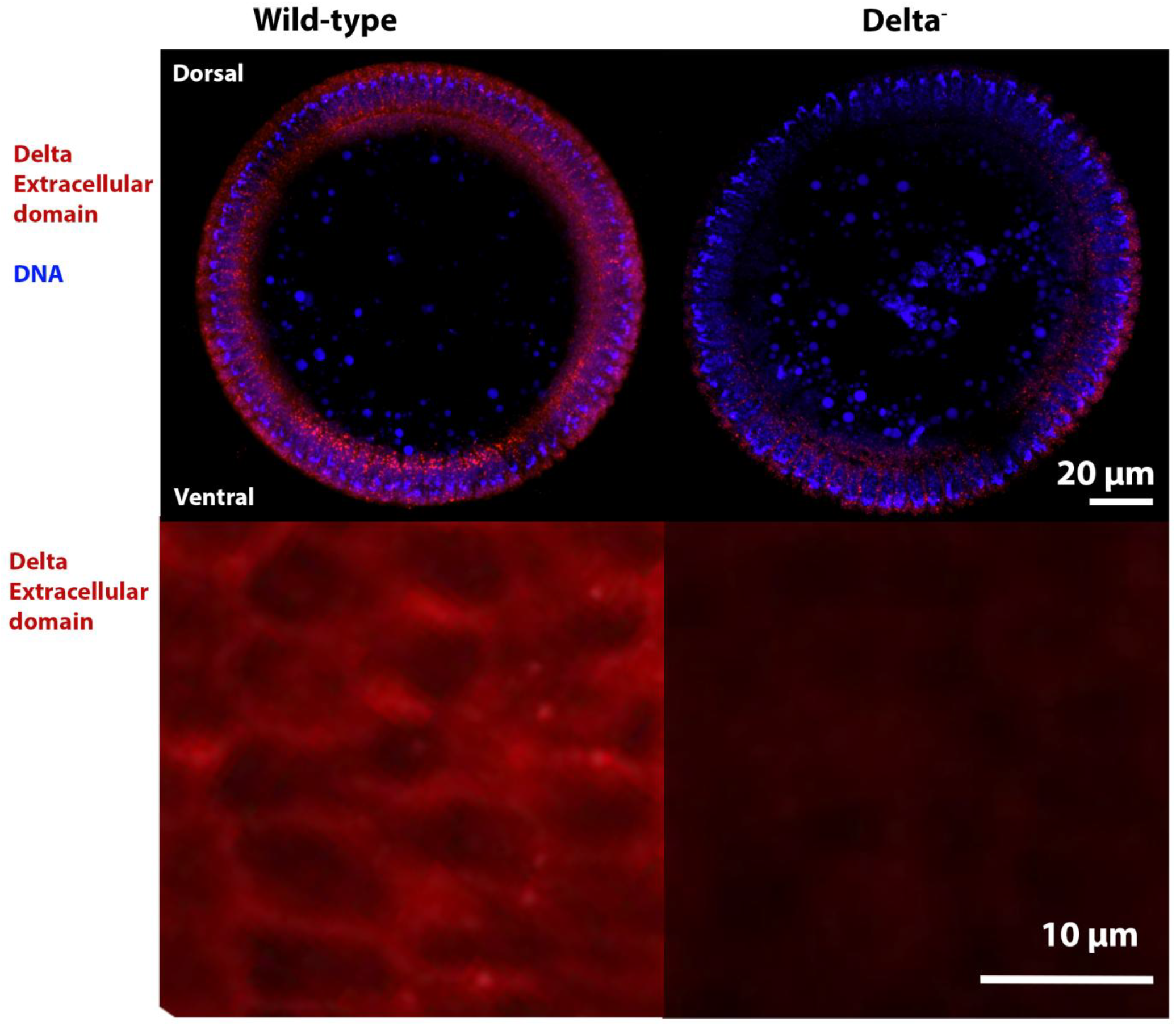
Heat-sensitive Delta mutation inhibits endocytosis of Delta when heat shocked at 32 °C, making it inaccessible for Notch extracellular domain and thereby inhibiting Notch signalling activation. A) Cross-sections of blastoderm wild-type and Delta^-^ embryos showing the decrease of Delta extracellular domain on cell membranes and intracellularly in Delta^-^ mutants compared wild-type embryos. B) Delta extracellular domain (red) expression in the ventral side of stage 6 wild-type and heat-shocked Delta^-^ mutant embryos. Delta extracellular domain is localized to cell membranes and inside the cells in wild-type embryos but is absent from most membranes of mutant embryos and no intracellular Delta is detected.

**Supplementary Table 1.**
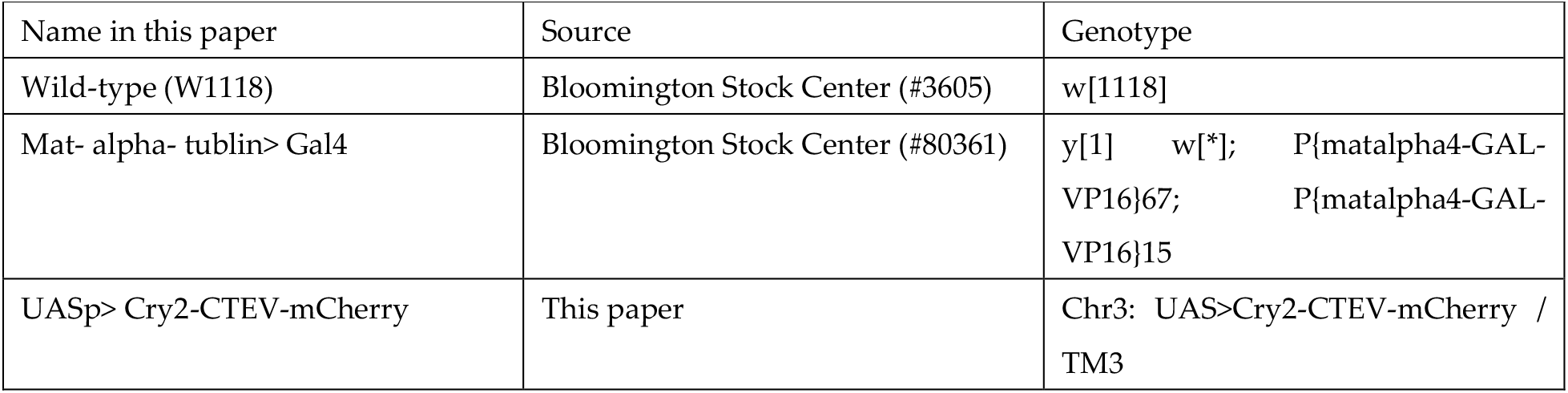
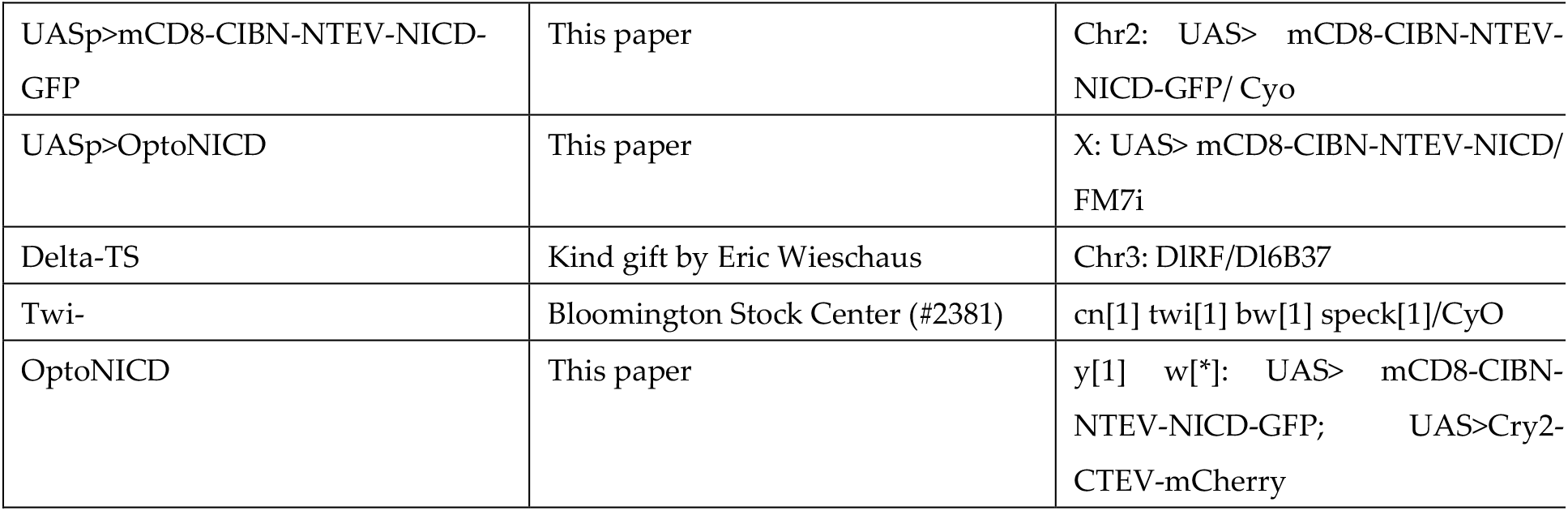
Flylines used and generated in this paper.

**Supplementary Table 2.**
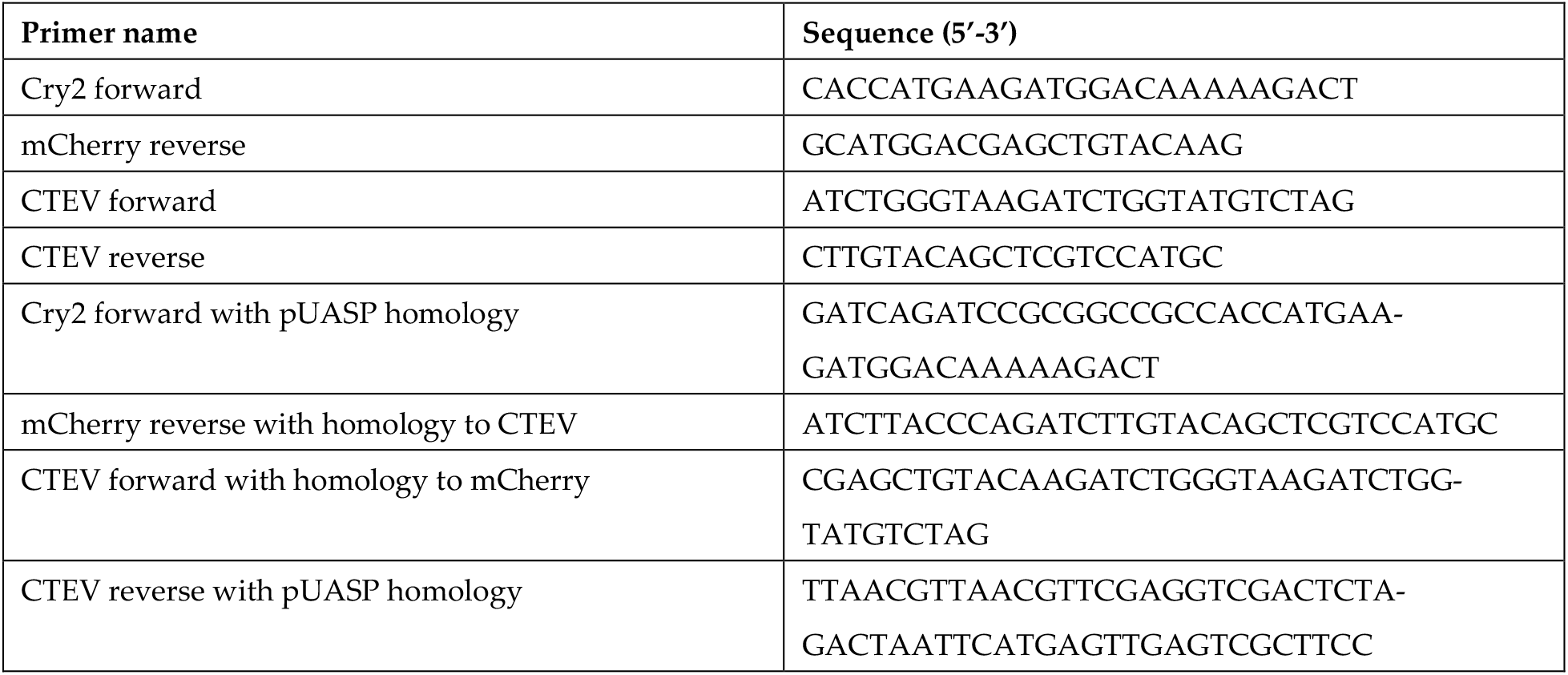
Primer sequences used for cloning.

**Supplementary Table 3.**
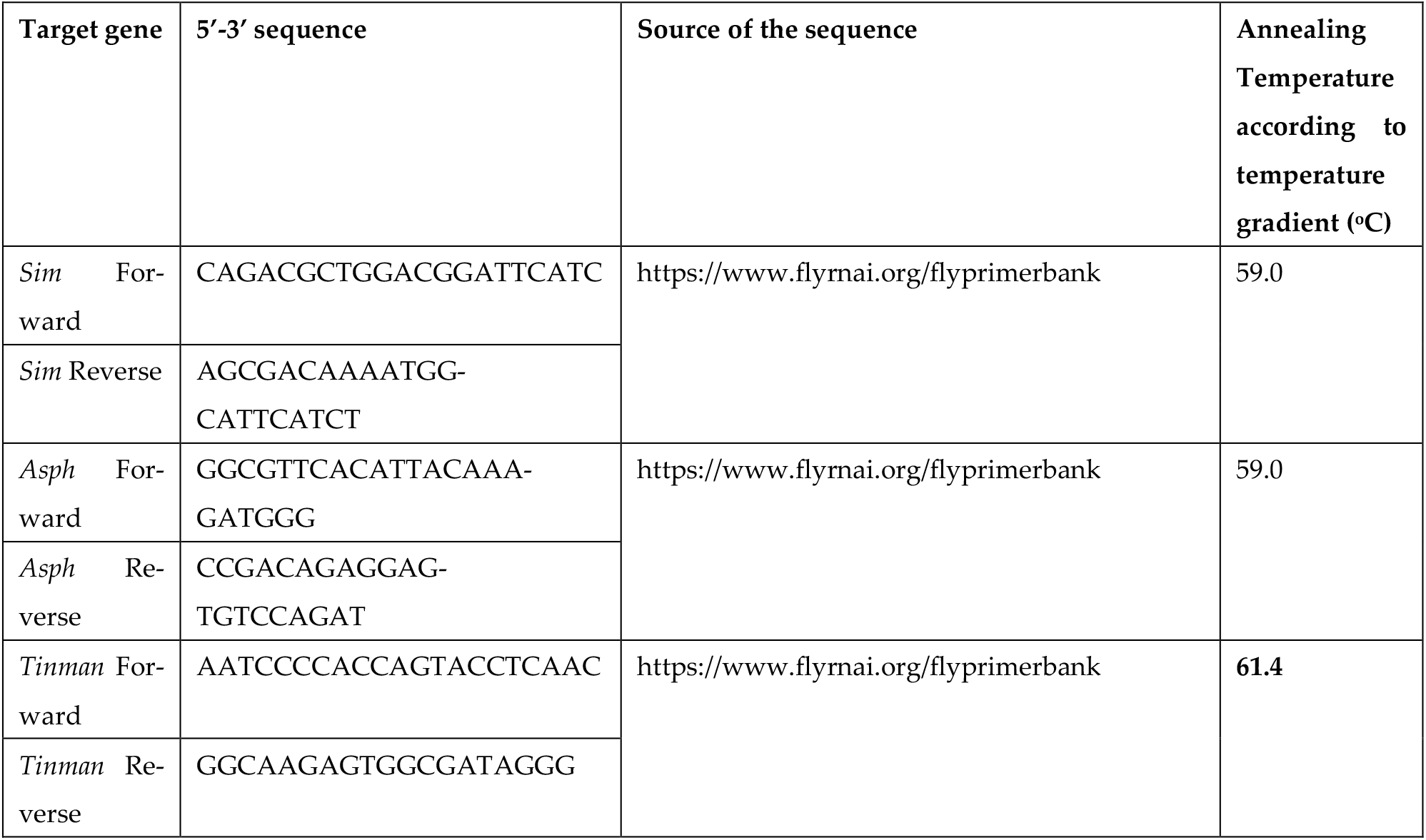
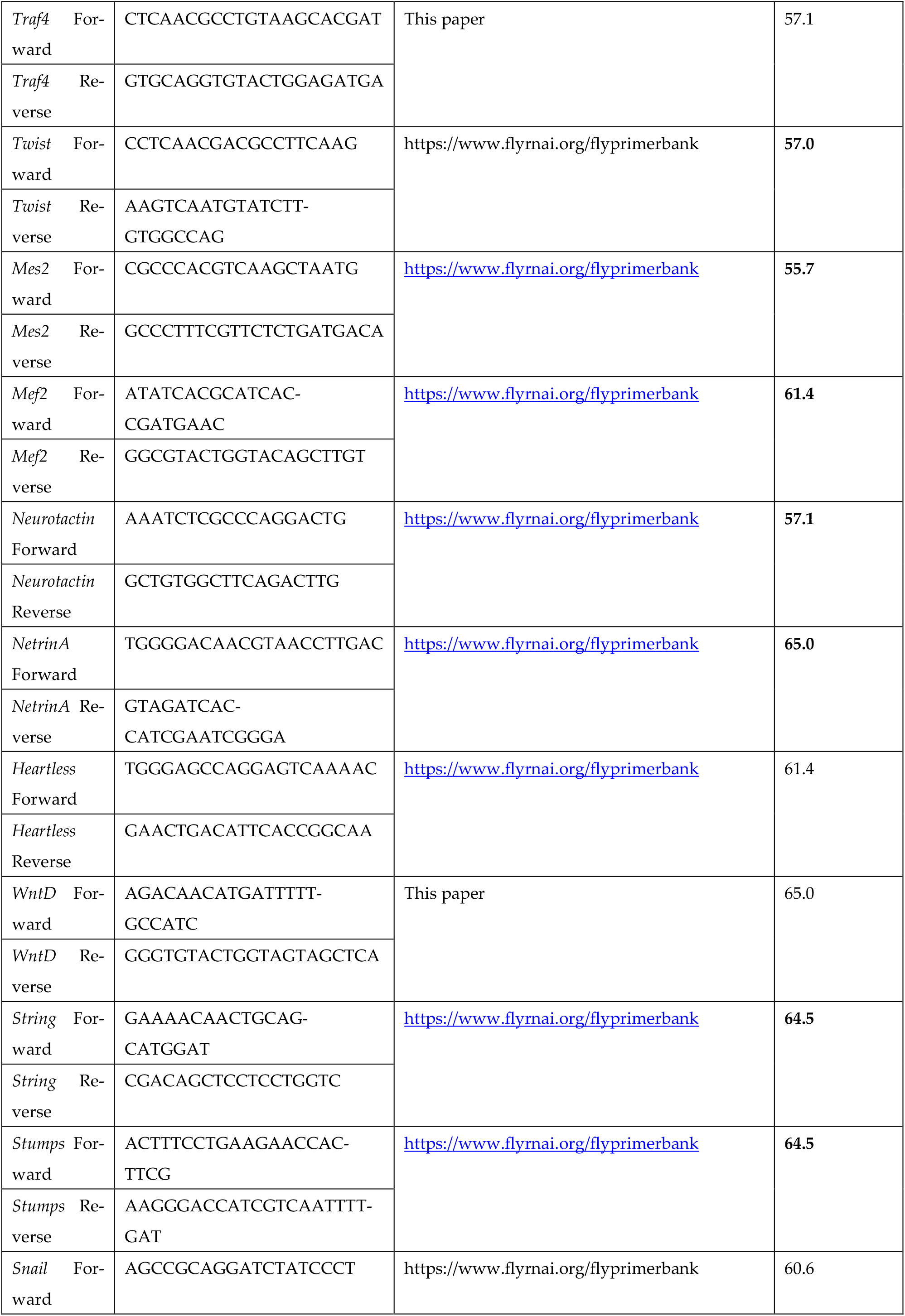
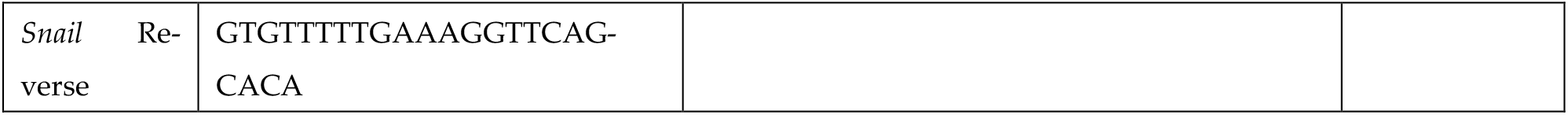
Primer sequences for qRT-PCR for mesodermal target genes.

**Supplementary table 4.**
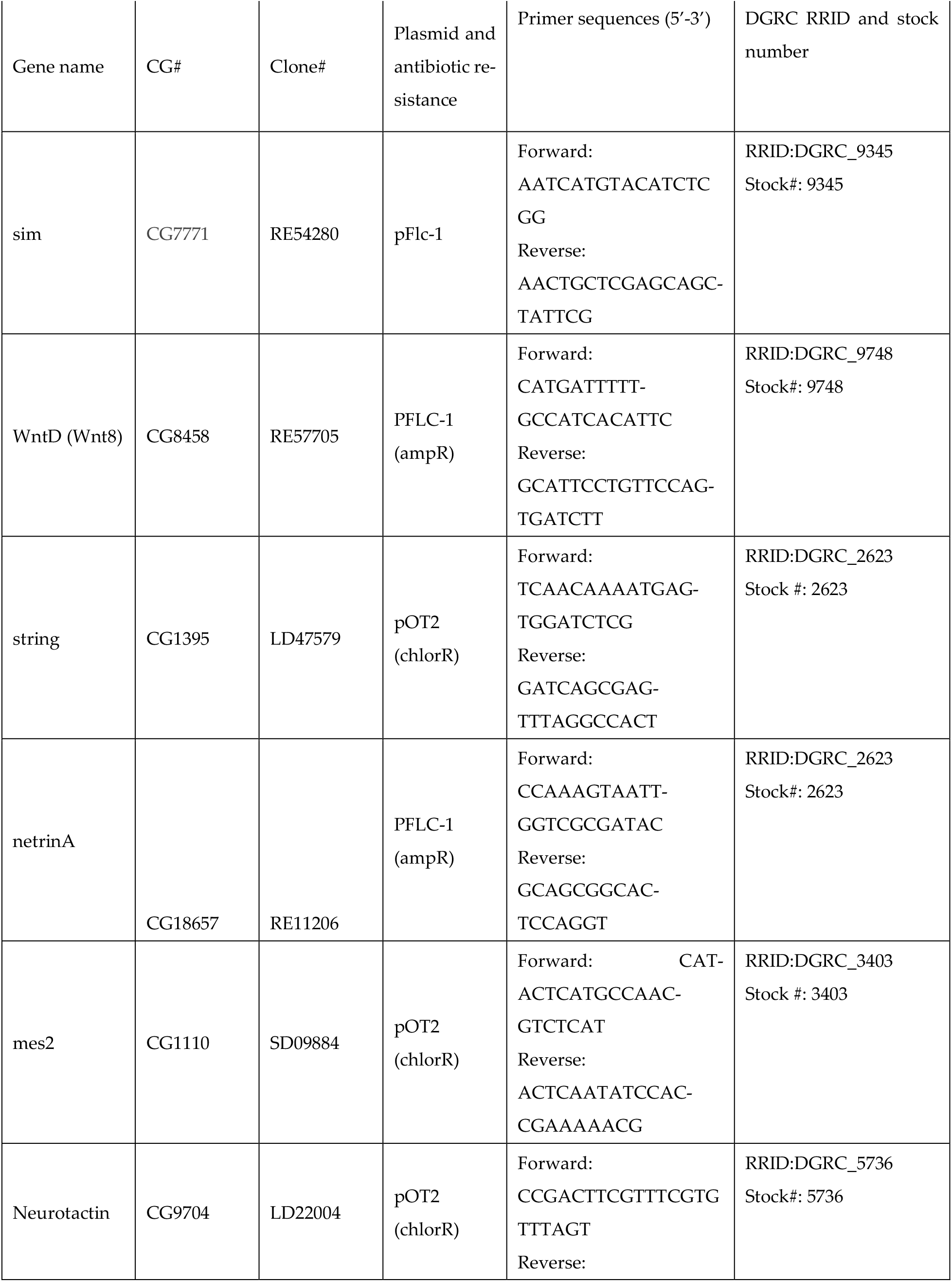
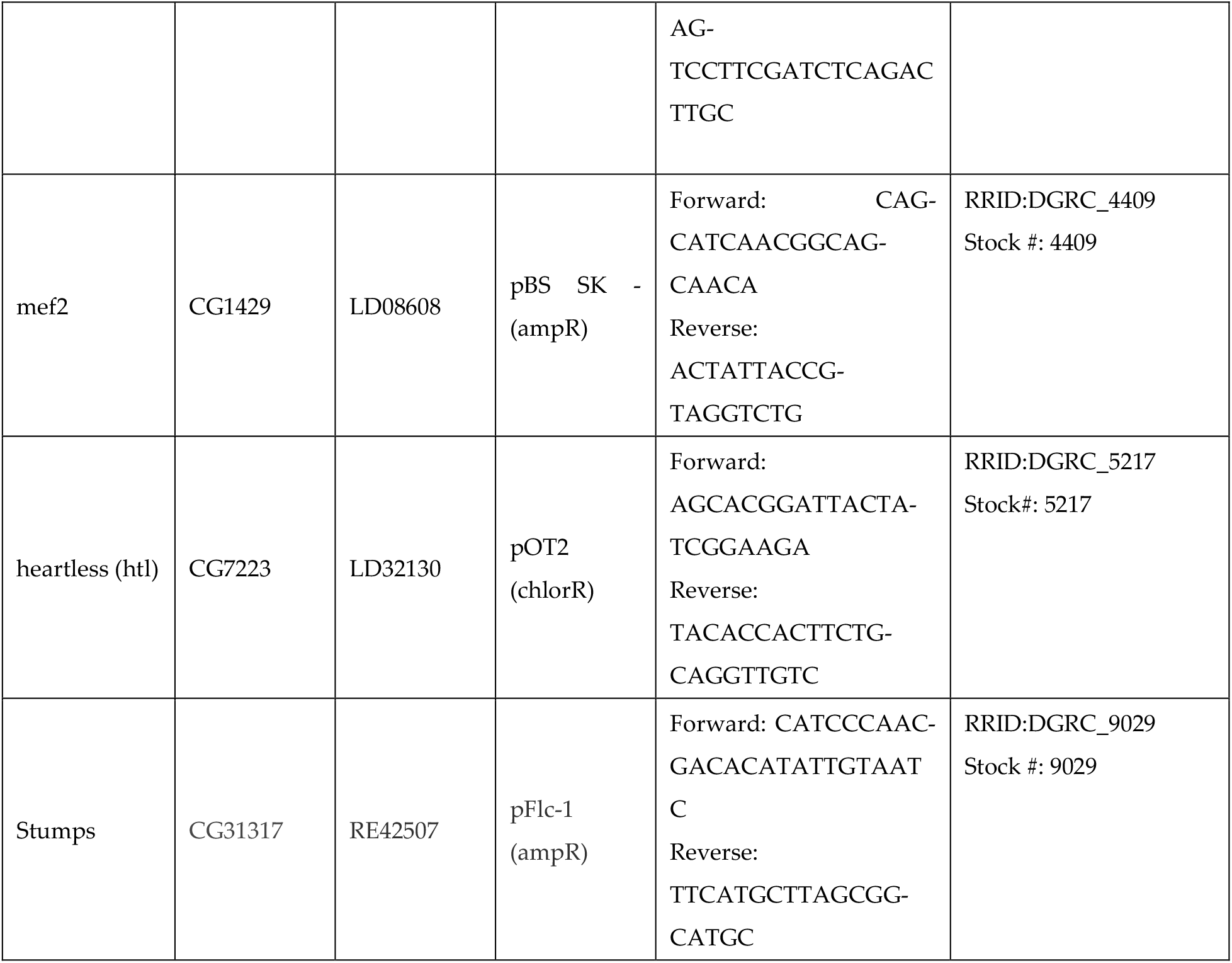
cDNA clones and primers used to amplify DNA templates to synthesize DIG and Biotin labeled antisense RNA probes for *in situ hybridizations*.

## Notes

### Competing Interest Statement

The authors have declared no competing interest.

### Summary of Updates

We made edits to the main text, clarifying some sections and improving the overall clarity of the paper. In addition, we included additional supplemental data (including additional controls) to support further the data represented in our main figures. We also updated the author list

